# A stromal-state dichotomy governs recalcitrance or remission in inflammatory bowel disease

**DOI:** 10.64898/2026.05.01.720931

**Authors:** Courtney Tindle, Harrison M. Penrose, Saptarshi Sinha, Madhubanti Mullick, Kennith Carpio-Perkins, Mason Hayashi, Sophia S. Carpinelli, Ella McLaren, Chia-Chian Hsieh, Kameron Zablan, Helen N. Le, Jennifer Neill, Gajanan D. Katkar, William J. Sandborn, Brigid S. Boland, Pradipta Ghosh

## Abstract

Inflammatory bowel diseases (IBD) remain a relapsing, treatment-refractory disorder marked by progressive tissue injury and inflammation despite expanding immune-targeted therapies. We established a prospective cohort integrating stromal biobanking, functional phenotyping, cross-cohort benchmarking, and outcome modeling to define disease-anchored cellular states. Colonic myofibroblasts from 34 individuals spanning health, ulcerative colitis, and Crohn’s disease resolved into **two dominant states**: inflammatory (**IMFs**) and quiescent (**QMFs**) myofibroblasts. QMF predominance forecasted remission, whereas IMF predominance increased the odds of worsening endoscopic severity despite therapy escalation during follow-up by ∼4.6-fold, linking early stromal biology to clinical outcomes. Unlike QMFs, IMFs exhibited a senescence-associated secretory phenotype that impaired epithelial stemness, barrier integrity, and innate immune fitness. State-guided prioritization identified EDNRB-antagonism as a high-confidence stromal intervention, reversing pathogenic phenotypes across orthogonal assays and species. Outcome simulation positioned stromal-state reversibility by EDNRB-antagonism as a precision axis, reducing odds of recalcitrance by ∼96.4% and reframing treatment resistance as a reversible stromal state.

**Graphic Abstract:** 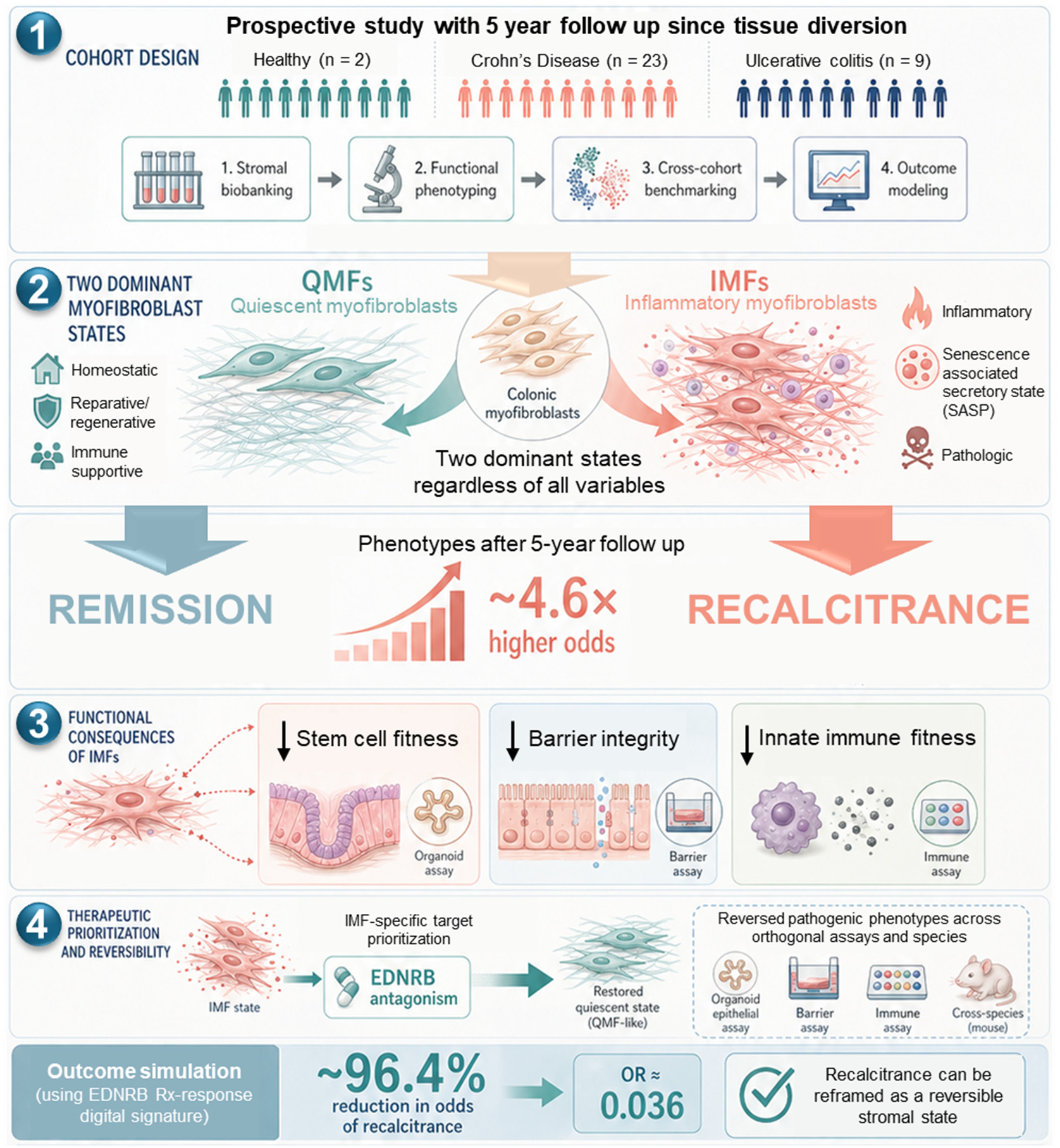

In this work, Tindle *et al.* identify reversible stromal states that govern remission vs. recalcitrant outcomes in IBD and nominate precision reprogramming of pathogenic myofibroblasts as a new therapeutic strategy.

## Introduction

Inflammatory bowel diseases (IBD) remain a relapsing, often recalcitrant disorder in which therapeutic resistance, treatment escalation, and progressive tissue injury persist despite expanding immune-targeted therapies^1–5^. A substantial proportion of patients fail to achieve durable remission and progress to develop strictures, fistulas, or require surgery^6–8^, with complication rates largely unchanged in the biologic era^9–11^. In Crohn’s disease (CD), this trajectory commonly culminates in fibrostenotic complications, including strictures, obstruction, fistulas^12^, and repeated resections^4^, whereas ulcerative colitis (UC) is characterized by persistent inflammation and mucosal dysfunction^13^. These patterns highlight a central unmet need: to define the cellular programs that drive disease progression and enable earlier, mechanism-based intervention.

Although IBD has traditionally been viewed through an immune-centric lens, emerging evidence implicates the stromal mesenchyme as a key regulator of progression and tissue remodeling^14–16^. Intestinal fibroblasts and myofibroblasts (**MF**s) reside at the mucosa–submucosa interface, where they integrate inflammatory cues with extracellular matrix (ECM) deposition, epithelial repair, and immune cell behavior, functioning as architectural hubs that couple active inflammation to structural damage^16–18^. Single-cell transcriptomic studies have deconstructed stromal heterogeneity to infinitesimal details, creating exhaustive catalogs of fibroblast subsets linked to epithelial support, immune modulation, therapy resistance, and fibrosis^18–29^ (see **Table 1**). However, these cross-sectional analyses, often performed on fibrotic tissues (end-stage disease), provide limited insight into precedent stromal states or how such states relate to future disease trajectory. Whether specific stromal programs actively shape clinical course remains unresolved; this is in part due to the lack of high-fidelity human new approach methodologies (NAMs) that preserve stromal biology in culture and inform longitudinal clinically relevant outcomes.

Here, we take a fundamentally different approach. We established a prospective, clinically anchored stromal biobank in which patient-derived cells were collected early and paired with long-term follow-up. This approach enables direct linkage of stromal states to disease trajectory. By integrating functional phenotyping with outcome modeling, we identify discrete stromal programs associated with progression and nominate actionable targets for intervention.

## Results

### Prospective human cohort reveals two stromal states in IBD

We established a prospective human cohort integrating endoscopic tissue diversion, primary stromal biobanking, and longitudinal follow-up (∼5 years) to anchor cellular states to clinical outcomes (**Fig. 1a; Supplementary Data 1**). To minimize selection bias, we implemented a label-free, marker-free outgrowth strategy in which mesenchymal cells migrating from biopsies were expanded as primary fibroblast/myofibroblast (MF) cultures (**Fig. 1b**). This approach mirrors widely used enzymatic methods^30,31^ while reducing potential perturbations associated with tissue homogenization or dissociation, including cellular stress, reduced viability, and altered surface protein integrity in sensitive cell populations^32^. Flow cytometry confirmed enrichment of CD45⁻ stromal cell populations and reproducible segregation into fibroblasts (CD90⁺αSMA⁻) and MFs (CD90⁺αSMA⁺), benchmarked against commercial controls (**Extended Data Fig. 1a, b**).

**Figure 1.**
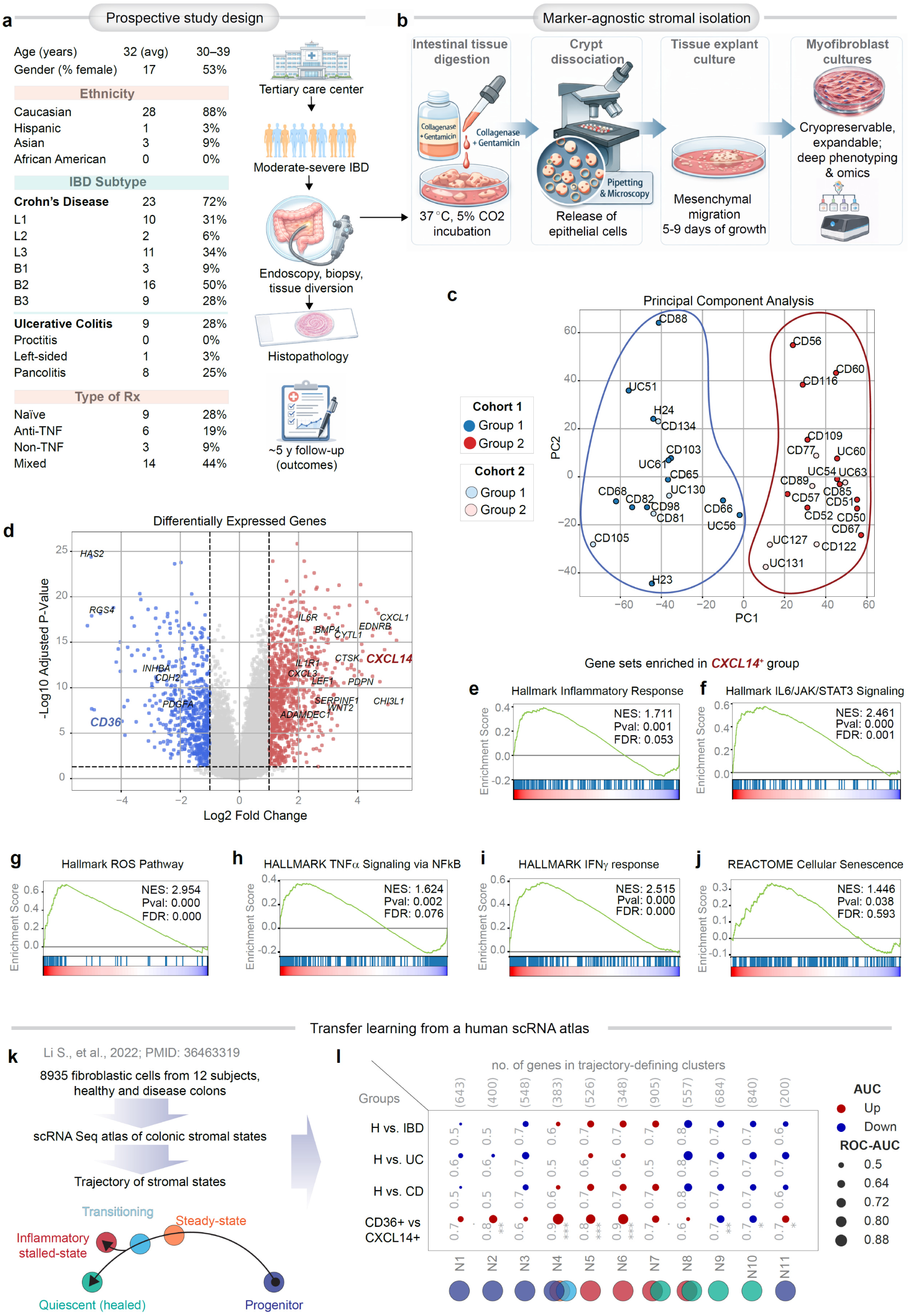
Prospective cohort framework and identification of stromal states in IBD. **a**, Patients with moderate-to-severe inflammatory bowel disease (IBD) were recruited from UC San Diego tertiary care center and underwent endoscopy with tissue diversion and biopsy. Clinical characteristics of the cohort at the time of tissue diversion are shown. Histopathology and longitudinal clinical outcomes (∼5-year follow-up beyond the time of diversion) were integrated with primary stromal cell biobanking. See **Supplementary Data 1** for detailed patient demographics. ‘Mixed’ indicates patients exposed to multiple biologics. **b**, Workflow for marker-agnostic isolation of intestinal stromal cells is shown. Mesenchymal cells migrating from tissue fragments over 5–9 days were expanded as primary myofibroblast cultures, enabling cryopreservation, phenotyping, and multi-omic analyses. See **Extended Data 1a-b** for marker-based flow cytometry. **c**, Principal component analysis of transcriptomes from primary myofibroblast cultures reveals two dominant stromal states (groups 1 and 2) that segregate across two independent cohorts, recruited over a 3-year period. **d**, Differential expression analysis identifies opposing transcriptional programs, CXCL14⁺ and CD36⁺. Key stromal-state marker genes are highlighted. See **Supplementary Data 2** for list of 224 genes. **e–j**, Gene set enrichment analysis plots show the most enriched pathways in the CXCL14⁺ state. See also **Extended Data 1c-f** for ontology analyses. **k,l**, Gene signatures of stromal states derived from the largest published human colonic scRNA-seq atlas (8,935 stromal cells; k) are used to map stromal trajectories in F/MF isolates of the current study. Dot plots (l) compare enrichment of stromal state signatures (columns; N1-11) in various comparator cohorts (rows); dot size reflects classification accuracy (AUC) and color denotes directionality (red, upregulated; blue, downregulated). No. of genes in each signature is indicated in parenthesis (top) and the list of genes in each signature is provided in **Supplementary Data 3**. *Statistics*: Enrichment statistics for pathways in e–j were computed using the gene set enrichment analysis algorithm, with normalized enrichment scores (NES) and corresponding false discovery rate (FDR) values reported. Statistical comparisons of composite gene score distributions between groups in panel l were performed using two-tailed, unpaired t-tests with Welch’s correction. *p* values are indicated in the panels; *P < 0.05, **P < 0.01, ***P < 0.001.

Three factors led us to focus on the colon as a clinically and biologically convergent site of IBD: (i) It is universally involved in UC and frequently affected in CD (∼60% of patients); while ∼half of those CD patients have synchronous involvement of the small intestine, the other half show disease that is limited to the colon^33^; (ii) Our prior work demonstrates that colonic epithelium in IBD segregates into two molecular subtypes, with surprising convergence of phenome, genome and transcriptome^34^, and that these epithelial programs forecast disease behavior across intestinal segments (such as complicated course of ileal CD)^35^; (iii) Others have shown that colonic stromal, but not their ileal counterparts, exhibit greater discriminatory power of complicated disease course across intestinal segments (fibrostenotic complication)^28^. This provided a rationale and high-yield substrate for stromal state discovery in the colon.

Our study design was intentional and forward-looking--by prospectively biobanking stromal cells with matched long-term outcomes, we sought to: (i) test whether stromal states track with objective disease trajectories (e.g., endoscopic severity, recalcitrant disease, etc.); (ii) identify actionable nodes capable of altering those trajectories, and (iii) establish biobanked stromal cultures as faithful human NAMs for therapeutic discovery. These goals distinguish this effort from prior foundational, but descriptive studies (see **Table 1**).

Unsupervised transcriptomic analysis revealed reproducible bifurcation of stromal identity. Principal component analysis segregated MFs into two near-equally represented clusters across two independently recruited cohorts (**Fig. 1c**), indicating a stable biological axis. Differential expression defined these states by opposing transcriptional programs centered on *CXCL14* and *CD36* (**Fig. 1d**), conventional markers of inflammatory vs quiescent states, respectively^22^ (see also **Table 1**). Notably, this dichotomy was invariant to diagnosis, demographics, and prior therapy exposure (all variables listed in **Supplementary Data 1**), revealing a binary stromal architecture that transcends conventional clinical stratification.

Functional annotation resolved the biological divergence of these states. *CXCL14*⁺ MFs were enriched for pathways linked to cytokine signaling, complement activation, and PI3K–Akt signaling (**Fig. 1e–j**; **Extended Data Fig. 1c,d**), consistent with an inflammatory, activated, tissue-remodeling phenotype. By contrast, *CD36*⁺ MFs were enriched for tissue morphogenesis, ECM organization, and focal adhesion programs (**Extended Data Fig. 1e,f**), consistent with structural maintenance and homeostasis.

To establish *in vivo* human relevance, we leveraged the largest available human colonic stromal single-cell atlas (8,935 cells; 12 subjects representing health and disease)^22^. Rather than segregating samples by health versus disease, stromal trajectories resolved along the CXCL14↔CD36 axis, with robust and consistent classification of key intermediate and activated states (N4-N6, N9-10, N-11; **Fig. 1k,l**). Compared to *CD36*⁺ MFs, *CXCL14*⁺ MFs were enriched in transitional (N5), inflammatory-stalled (N6-7), and progenitor-like (N11) states, with concomitant suppression of quiescent programs (N9-10) (**Fig. 1l**). These observations motivated functional nomenclature: *CXCL14*⁺ inflammatory MFs (IMFs) and *CD36*⁺ quiescent MFs (QMFs).

Together, these data define a conserved, binary stromal architecture in the colon, IMF-dominant and QMF-dominant states, that is independent of traditional clinical categories and provides a mechanistic foundation to link stromal state composition to disease trajectory in IBD.

### IMF predominance anchors stromal biology to risk of recalcitrance in IBD

Building on the identification of a binary stromal architecture, we next asked whether these states represent a conserved disease axis across independent datasets and whether they carry clinical consequence. To test this, we performed cross-cohort benchmarking against 12 independent single-cell RNA sequencing IBD stromal signatures spanning UC and CD, derived from both large atlases and focused mechanistic studies, including perianal fistulizing disease (**Table 2**)^12,18,20,21,23,25,27–29,36–45^. This analysis integrated 549 human samples across diverse clinical phenotypes and experimental platforms (**Fig. 2a**). Despite differences in patient demographics, tissue handling, sequencing strategies, and analytical frameworks across studies, comparative enrichment analysis demonstrated robust and consistent classification performance of these signatures. High area-under-the-curve (AUC) values across the majority of stromal signatures confirmed that disease-associated states prioritized in prior work (see **Table 2**) maps coherently onto the disease (UC, CD, or IBD) derived MF biobank established here (**Fig. 2b-** *top three rows*, red). Most of those disease-enriched states mapped onto *CXCL14*⁺ IMFs, with improved significance (**Fig. 2b-** *bottom row*, red). Some of the homeostatic healthy states were depleted in disease derived MFs (**Fig. 2b**- *top 3 rows*, blue), as well as in IMFs (**Fig. 2b**- *bottom row*, blue). Together, these data define a conserved, binary stromal architecture that provides a foundation to link stromal state composition to disease course in IBD. This convergence across independent human datasets establishes that the IMF–QMF axis represents a conserved and biologically dominant stromal architecture in IBD, providing a rigorous foundation to interrogate its relationship to disease trajectory.

**Figure 2.**
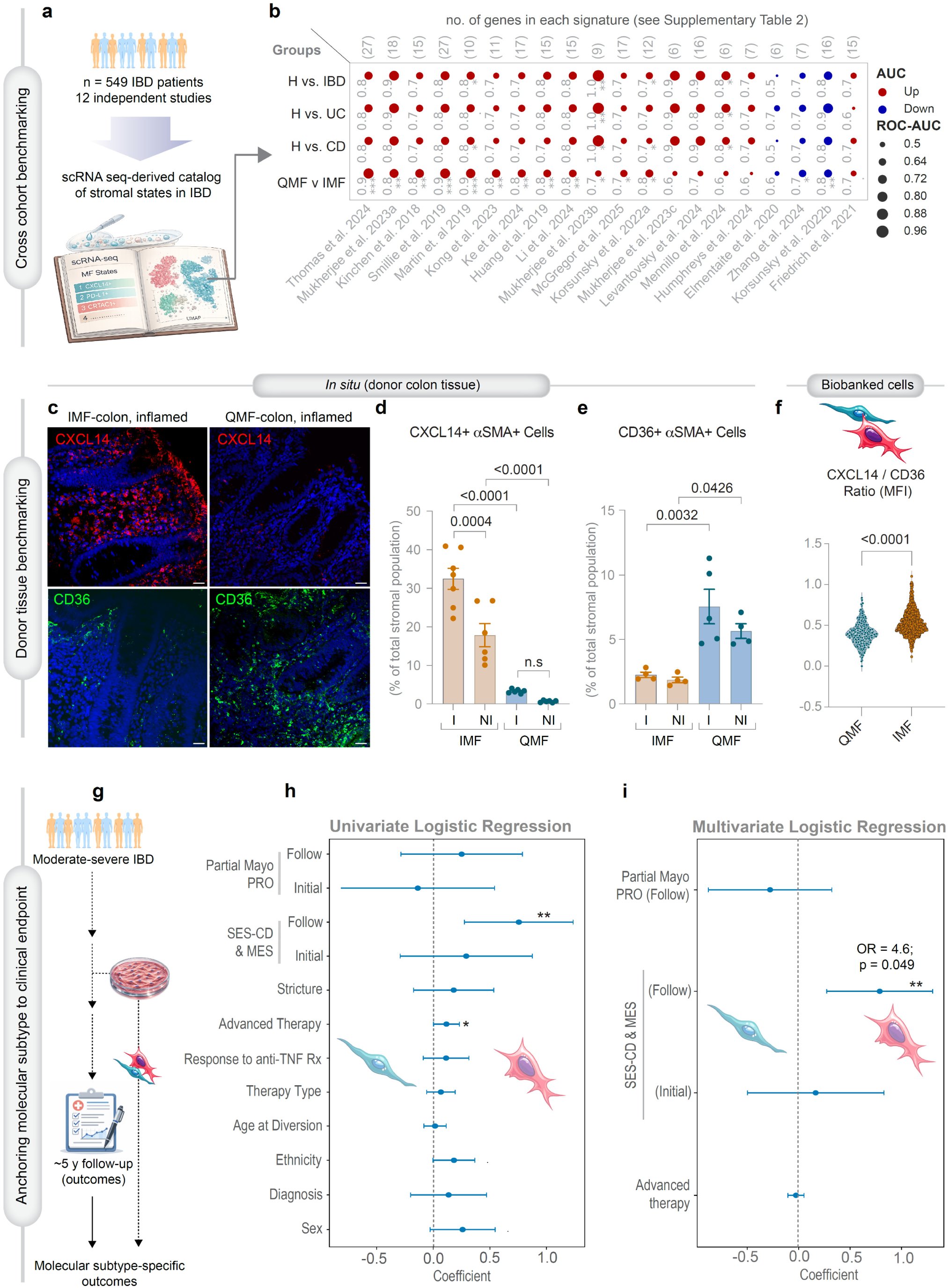
IMF predominance anchors stromal biology to clinical risk of progression in IBD. **a-b, Cross-cohort benchmarking of stromal states**. Gene signatures defining IBD-associated stromal states in 12 independent studies (**a**; listed in **Table 2**) were analyzed in the F/MF isolates of the current study. Dot plots (**b**) compare enrichment of IBD-associated stromal state signatures (columns; **Table 2**) in various comparator cohorts (rows); dot size reflects classification accuracy (AUC) and color denotes directionality (red, upregulated; blue, downregulated). No. of genes in each signature is indicated in parenthesis (top). Statistical comparisons of composite gene score distributions between groups were performed by two-tailed, unpaired t-tests with Welch’s correction; *P* values indicated in panels; P < 0.05 (*); <0.01 (**); <0.001 (***). See **Table 2** for detailed p values associated with each gene signatures. **c–e, Donor tissue benchmarking at time of diversion.** Representative confocal immunofluorescence images (**c**) of colon tissue sections obtained at the time of diversion, stained for CXCL14⁺ IMFs (red) and CD36⁺ QMFs (green) with nuclei counterstained (blue; DAPI). Scale bar, 20 µm. See **Extended Data 2-3** for representative images from non-inflamed regions of the same patients. Quantification (**d-e**) was performed across 4-7 fields of view (FOVs) per section, with stromal populations identified by co-localization with αSMA⁺ cells. Bars indicate mean ± S.E.M. I, inflamed; NI, non-inflamed. **f, Biobanked stromal cell phenotyping.** Primary myofibroblasts derived from patient tissues were expanded *ex vivo* and quantified for CXCL14/CD36 expression ratios by immunostaining and image-based quantification. See **Extended Data 4** for representative confocal immunofluorescence images. **g, Schematic of clinical anchoring framework**. Stromal cell states at diversion were linked prospectively to longitudinal clinical outcomes, with approximately five years of follow-up. **h,i, Association of stromal states with clinical outcomes.** Forest plots show coefficients from univariate (**h**) and multivariate (**i**) ordinary least squares models evaluating the relationship between stromal state predominance (base variable) and clinically relevant patient characteristics and endpoints. Multivariate models (**i**) adjusted for clinical covariates demonstrate that IMF-state signature at the time of tissue diversion predicts persistent endoscopic inflammation at follow-up, as measured by worsening endoscopic scores [Simple Endoscopic Score for Crohn’s Disease (SES-CD) and Mayo Endoscopic Subscores (MES) for ulcerative colitis, see ***Methods***] (OR = 4.6, 95% CI 0.89–23.8, Fisher exact p = 0.049). See **Supplementary Data 1** for detailed patient information. PRO, patient reported outcomes. *Statistics*: Statistical comparisons in **d-e** were performed using one-way ANOVA. In **f**, each point represents single cell analyses in culture; distributions are shown as violin plots with median and interquartile ranges. Differences were assessed using a two-sided, unpaired t-test. In **h-i**, Ordinary least squares models report coefficients with corresponding significance levels. Points represent regression coefficients with 95% confidence intervals; significance is indicated in panels. Two tailed *p* values indicated in panels; P < 0.05 (*); <0.01 (**).

We then anchored these states directly to donor tissue at the time of patient recruitment and biobank creation. Confocal imaging of colonic sections obtained at diversion revealed spatially distinct stromal compartments marked by CXCL14⁺ IMFs and CD36⁺ QMFs (**Fig. 2c, Extended Data Fig. 2-3**). Quantification across multiple fields of view demonstrated marked inter-patient dichotomy, with donor tissues mirroring IMF or QMF predominance *in vivo* (**Fig. 2d,e**), as did their corresponding cultures *ex vivo* (**Fig. 2f; Extended Data Fig. 4**). Importantly, these patterns were not stochastic: spatial analyses across inflamed and non-inflamed regions confirmed that CXCL14⁺αSMA⁺ stromal compartments selectively expand within inflamed niches, whereas CD36⁺αSMA⁺ cells are relatively enriched in non-inflamed or reparative regions (**Fig. 2c**). This establishes that transcriptionally defined states in cultures are preserved *in situ* and organized within the tissue microenvironment. That patient-specific IMF versus QMF predominance was faithfully preserved in culture indicates that these states are stable, intrinsic properties rather than transient, context-dependent artifacts. This stability supports their use as tractable human NAMs for mechanistic interrogation.

We next anchored stromal states to clinical outcomes. Using a prospective framework (**Fig. 2g**), stromal state predominance at diversion (IMF vs. QMF, as a binary variable) was assessed in relationship to longitudinal disease trajectories over ∼5 years. IMF predominance associated strongly with adverse outcomes: In univariate models, this tracked with objective measures of disease severity, including worse endoscopic scores and need for therapy escalation during follow-up (**Fig. 2h**). In multivariate models adjusting for clinical covariates, IMF predominance remained an independent predictor of persistent endoscopic inflammation despite therapy escalation, as measured by worsening endoscopic scores at follow-up (OR = 4.6, 95% CI 0.89–23.8, p = 0.049; **Fig. 2i)** (see ***Methods*** for clinical outcomes analysis).

Together, these data move stromal states from descriptive taxonomy to clinical axis. IMF predominance emerges as a stable, spatially encoded, and prospectively validated actionable dimension of recalcitrant IBD, motivating efforts to therapeutically reprogram this axis.

### IMFs define a senescent, metabolically active, immunotolerant stromal niche

We next defined the phenotypic programs underlying IMF versus QMF states and anchored these behaviors to the transcriptional signatures identified in **Fig 1**. Donor-derived stromal lines representing exemplar extremes of the IMF↔QMF continuum were subjected to orthogonal phenotyping across early passages to assess stability and functional divergence (**Fig. 3a**).

**Figure 3.**
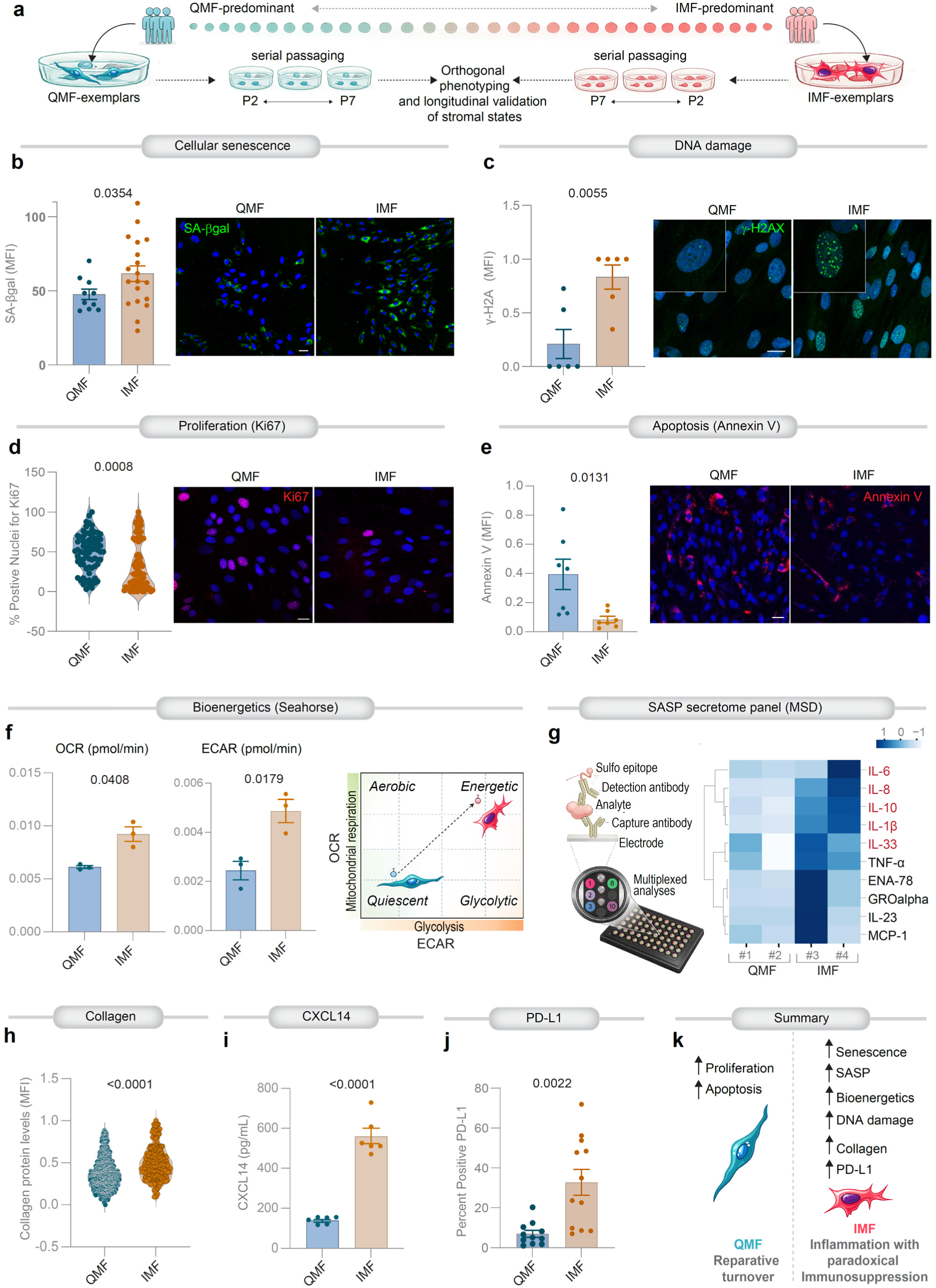
Inflammatory MFs encode a senescence-associated, paradoxically immunotolerant stromal niche. **a, Experimental design**. Donor-derived stromal cell lines were ranked along a continuum from QMF-predominant to IMF-predominant states. The most divergent lines from each pole were selected as exemplars and subjected to serial passaging (limited to < P7) and orthogonal phenotyping to assess longitudinal stability of stromal states. **b**, Quantification of Senescence-associated β-galactosidase (SA-β-gal)–positive cells by flow cytometry are shown alongside representative fluorescence images (green, SA-β-gal; blue, nuclei). Scale bar, 20 µm. Each point on the bar graph represents an independent donor-derived culture; n = 7 unique IMF and 4 unique QMF subjects. 2-3 unique passages/subject. Bars represent mean ± S.E.M. Statistical significance was assessed using two-sided, unpaired t-tests. **c**, Quantification of mean fluorescence intensity (MFI) from γ-H2AX staining, as determined by flow cytometry is shown with representative fluorescence images (green, foci of DNA double-stranded breaks; blue, nuclei). Scale bar, 20 µm. Each point represents an independent donor-derived culture; n = 3 unique IMF and 3 unique QMF subjects; 1-3 unique passages/subject. Error bars represent mean ± S.E.M. Statistical significance was assessed using two-sided, unpaired t-tests. **d**, Violin plots show the distribution of Ki67-positive nuclei across biological replicates, with representative microscopy images (red, actively dividing stained cells; blue, nuclei). Scale bar, 20 µm. Each point represents an independent donor-derived culture; n = 3 unique IMF and 3 unique QMF subjects; 2-4 unique passages/subject, with 10 high-power fields/technical repeat. Error bars represent mean ± S.E.M. Statistical significance was assessed using two-sided, unpaired t-tests. **e**, Quantification of Annexin V MFI is shown with representative fluorescence images (red, apoptotic cell; blue, nuclei). Scale bar, 20 µm. Each point represents an independent donor-derived culture; n = 2 unique IMF and 2 unique QMF subjects; 4 unique passages/subject, with 3 high-power fields/technical repeat. Error bars represent mean ± S.E.M. Statistical significance was assessed using two-sided, unpaired t-tests. **f**, Quantification of oxygen consumption rate (OCR; left) and extracellular acidification rate (ECAR; middle) is shown. See also **Extended data 5a-b** for plots of seahorse real-time metabolic flux analysis. Conceptual metabolic phenotyping plots OCR against ECAR, illustrating state transition between QMF and IMF. Each point represents an independent donor-derived culture; n = 4 unique IMF and 4 unique QMF subjects; 3 unique passages/subject. Error bars represent mean ± S.E.M. Statistical significance was assessed using two-sided, unpaired t-tests. **g**, Multiplex electrochemiluminescence immunoassay (MSD platform; left) used to quantify cytokines and chemokines secreted by stromal cells. Heatmap (right) shows a panel of SASP-related inflammatory mediators. Supernatants pooled from three unique passages of each patient is shown. Highlighted (red) indicate consistently elevated cytokines in IMF (see **Extended data 5c**). **h–j**, Multiple orthogonal assays assess stromal effector programs, e.g., quantification of collagen (**h**), secretion of the IMF-defining chemokine (CXCL14; **i**), and expression of the immune checkpoint ligand PD-L1 (**j**). In h, each point represents single cell analyses in culture; distributions are shown as violin plots with median and interquartile ranges. See **Extended data 5d** for morphology and collagen-stained fields. In i, each point represents an independent donor-derived culture; n = 2 unique IMF and 2 unique QMF subjects; 3 unique passages/subject. In j, each point represents an independent donor-derived culture; n = 4 unique IMF and 4 unique QMF subjects; 2-5 unique passages/subject. **k**, Summary of findings that define contrasting stromal programs. Senescent IMFs promote inflammation while simultaneously establishing a locally immunotolerant niche. Reparative QMFs show enhanced turnover. *Statistics*: For all panels, points represent independent donor-derived cultures unless otherwise indicated. Representative microscopy images are shown from multiple biological replicates with multiple fields of view per condition. Statistical comparisons were performed using two-sided, unpaired t-tests; exact *P* values are shown.

Consistent with enrichment of inflammatory and stress-response programs (**Fig. 1e–j**), IMFs exhibited hallmark features of cellular senescence. Relative to QMFs, IMFs showed increased senescence (elevated β-galactosidase activity; **Fig. 3b**) and accumulation of DNA damage (abundance of γ-H2AX foci; **Fig. 3c**). This was accompanied by reduced proliferative capacity (fewer Ki67⁺ nuclei, **Fig. 3d**) and diminished apoptosis (lower Annexin V staining; **Fig. 3e**). Together, these features define a senescent stromal population that resists turnover and persists.

We next interrogated metabolic programs, guided by enrichment of PI3K–Akt and stress-response pathways in IMFs (**Fig. 1e–j**). Seahorse-based flux analyses revealed that IMFs are metabolically hyperactive, with increased oxidative phosphorylation and glycolytic flux relative to QMFs (**Fig. 3f**; **Extended Data Fig. 5a,b**). This dual activation supports a sustained, energetically competent effector state, consistent with enrichment of transitional and progenitor-like populations within the IMF axis (**Fig 1l**), rather than terminal quiescence.

Secretome profiling linked transcriptional identity to functional output. Multiplex cytokine analysis demonstrated robust induction of senescence-associated secretory phenotype (SASP) mediators in IMFs, including IL-6, IL-8, IL-1β, MCP-1, TNF-α, and CXCL1 (**Fig. 3g; Extended Data Fig. 5c**). CXCL14, a defining transcriptomic marker (**Fig. 1d**), was consistently elevated at the protein level (**Fig. 3i**), underscoring tight coupling between gene expression and effector function.

Beyond inflammatory signaling, IMFs exhibited coordinated programs of matrix remodeling and immune modulation. IMF cultures showed increased collagen production and cytoskeletal organization (**Fig. 3h; Extended Data Fig. 5d**), consistent with enrichment of ECM and tissue-remodeling pathways (**Fig. 1e–j**). Notably, IMFs upregulated the immune checkpoint ligand Programmed death-ligand 1 (PD-L1; **Fig. 3j**), a feature conserved across diverse tissues (gut, lung and liver) that is associated with stromal activation, persistence and resistance to immune-mediated clearance^31,46–51^.

These findings reveal a paradoxical phenotype: IMFs couple a pro-inflammatory secretome with establishment of a locally immunotolerant niche. By contrast, QMFs displayed a complementary phenotype characterized by lower senescence burden, preserved proliferative capacity, reduced metabolic activity, and minimal SASP output, consistent with their enrichment for morphogenetic and homeostatic pathways (**Extended Data Fig. 1e,f**).

Taken together, these data functionally resolve the IMF–QMF axis. IMFs represent a senescence-associated, metabolically active, and immunomodulatory stromal state that integrates inflammatory signaling with local immune suppression, whereas QMFs maintain a reparative, homeostatic program.

### IMF paracrine signaling disrupts epithelial integrity and impairs innate immune function

We next asked whether the IMF–QMF stromal programs translate into functional consequences for neighboring epithelial and immune compartments. We interrogated stromal–epithelial and stromal–myeloid crosstalk using conditioned media from donor-derived QMF and IMF cultures (**Fig. 4a,d,g**).

**Figure 4.**
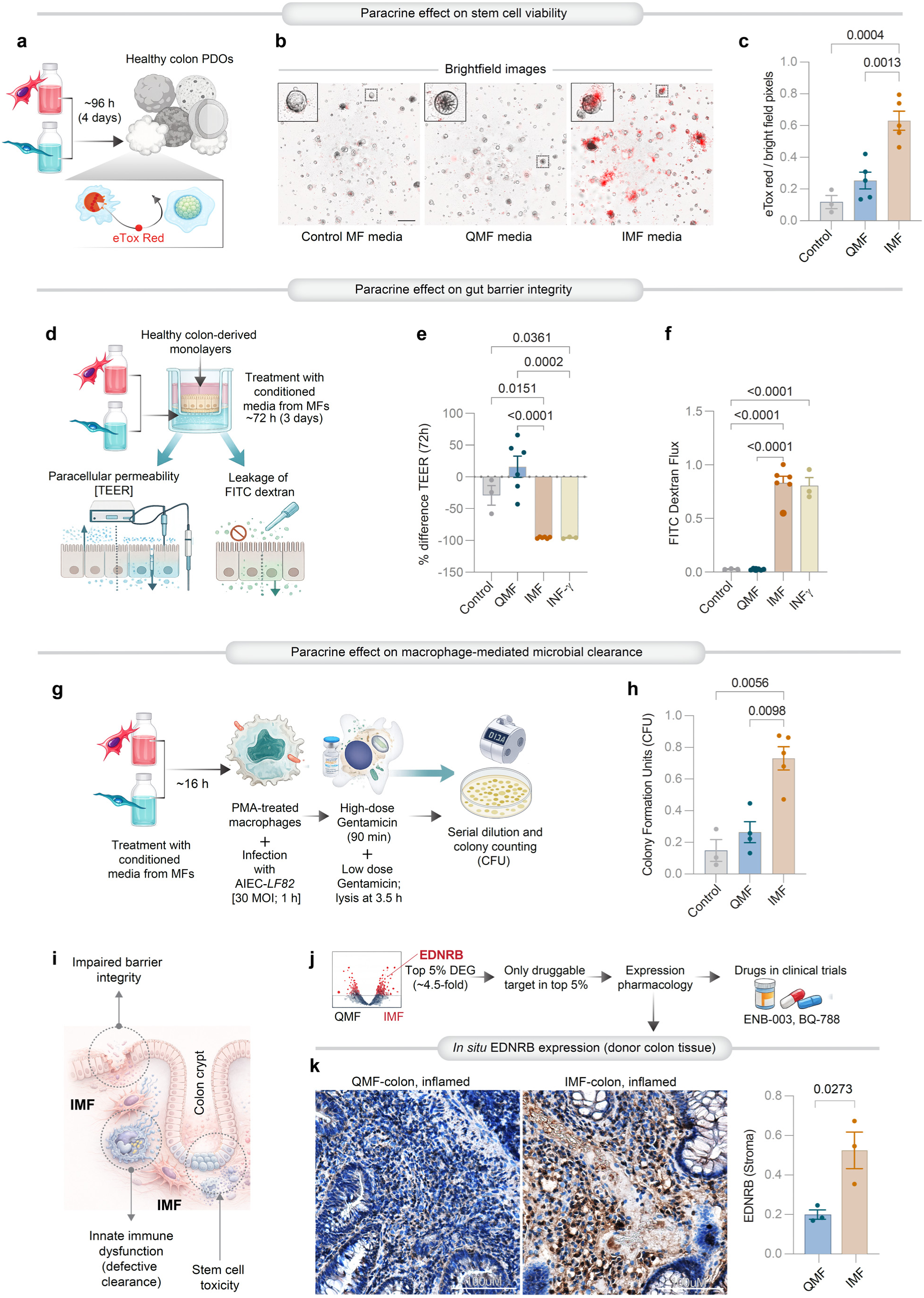
Stromal paracrine crosstalk impairs epithelial and myeloid fitness. **a–c**, Schematic of workflow (**a**) shows key steps of organoid viability assays, as assessed using the cell-impermeant dye eTox Red, which labels dying cells. Representative brightfield images (**b**) of organoids subjected to culture in the presence of various conditioned media. Insets show representative organoids at higher magnification. Quantification (**c**) of eTox Red–positive pixels normalized to brightfield pixels. Each point represents conditioned media from independent donor-derived cultures; n = 2 unique IMF and 2 unique QMF subjects; 2-3 unique passages/subject. Representative images are derived from 2-3 fields of view per condition per replicate. Scale bar, 0.5 mm. **d–f**, Schematic of workflow (**d**) shows key steps of two orthogonal assays of barrier function, transepithelial electrical resistance (TEER) and FITC-dextran permeability assays. Quantification (**e**) of % change in TEER after treatment with conditioned media from control MFs, QMFs, or IMFs, compared with interferon-γ (IFNγ)–treated positive control. See **Extended Data 6a-b** for time-course measurements (0–72 h). Quantification (**f**) of FITC-dextran flux across epithelial monolayers, assessed under the same conditions as e. Results represent findings of conditioned media from independent donor-derived cultures; n = 2 unique IMF and 2 unique QMF; 3 unique passages/subject. **g–h**, Schematic of workflow (**g**) shows key steps of microbial clearance assays on PMA-differentiated THP1-derived macrophages. Quantification (**h**) of bacterial CFUs recovered from macrophages, showing microbes that survive innate immune clearance. Each point represents conditioned media from independent donor-derived cultures; n = 2 unique IMF and 2 unique QMF subjects; 2-3 unique passages/subject. **i**, Summary of findings: IMF-dominated stromal states exert paracrine effects of IMFs in promoting epithelial injury and innate immune dysfunction. **j**, Systematic prioritization identifies EDNRB as the only clinically actionable target within the IMF program. **k**, Representative immunohistochemistry (IHC; left) images and quantification of inflamed colonic regions from IMF- or QMF-predominant tissues stained for EDNRB. Scale bar 100 µm. Quantification (right) of stromal EDRNB was performed across 3 fields of view (FOVs) per section. Bars indicate mean ± S.E.M. See **Extended Data 7** for multiple FOVs and secondary controls. *Statistics*: Data represents unique subjects, and technical repeats, as indicated. Bars denote mean ± S.E.M. Statistical comparisons were performed using one way ANOVA or two-sided, unpaired t-tests with exact *P* values shown.

Exposure of healthy human colonic organoids to IMF-conditioned media resulted in marked loss of viability, with increased uptake of cell-impermeant dye (eTox Red), indicating epithelial injury (**Fig. 4a–c**). By contrast, QMF-conditioned media preserved organoid integrity, resembling control conditions. These findings establish that IMF-derived secretomes are sufficient to drive epithelial damage. Given that 3D patient-derived organoid (PDO) cultures are enriched for stem-like and transient amplifying cells, these findings align with prior observations of stem cell dysfunction^52,53^, telomere injury^54^, and increased apoptosis^55^ seen in IBD PDOs by others and us^34^, positioning IMF-derived signals as direct disruptors of the regenerative compartment.

We next assessed barrier function using complementary assays on healthy human colonic PDO-derived monolayers. IMF-conditioned media triggered a pronounced reduction in transepithelial electrical resistance (TEER), approaching levels seen with the known disruptor interferon-γ^56–58^ (**Fig. 4d,e; Extended Data Fig. 6a,b**), and increased paracellular permeability, as measured by FITC-dextran flux (**Fig. 4f**). By contrast, QMF-conditioned media had no discernible effect in both assays. These data mechanistically link IMF-derived factors to epithelial barrier failure, mirroring defects reported in IBD epithelial monolayers by others^59,60^ and us^34^.

We then examined effects on innate immune function. To model the dysbiotic IBD lumen, we used live cultures of the pathogenic adherent invasive *Escherichia coli* strain-*LF82* (AIEC*-LF82*), because it was isolated from CD patients^61^. Macrophages exposed to IMF-conditioned media exhibited impaired AIEC*-LF82* clearance, with increased recovery of viable bacteria following infection (**Fig. 4g,h**), whereas QMF-conditioned media had a negligible effect.

Together, these data establish stromal state composition as a driver of epithelial and innate immune dysfunction (**Fig. 4i**), two core features of recalcitrant IBD^62–65^. IMF-dominated niches simultaneously induce epithelial injury and barrier breakdown while impairing microbial clearance, positioning IMFs as central orchestrators of a self-reinforcing cycle of tissue damage and immune failure.

### EDNRB antagonism reverses IMF-driven pathology and restores epithelial and immune fitness

Because of their potential as drivers of recalcitrant IBD, we next sought to therapeutically target the IMF stromal axis. Target prioritization was guided by the transcriptional architecture defined in **Fig. 1** (**Fig. 4j**). Among the most upregulated genes in IMFs (top 5% differentially expressed genes; ∼4.5-fold change; **Fig. 1d**; **Supplementary Data 2**), endothelin receptor type B (EDNRB) emerged as a uniquely actionable node. It was the only druggable target within this tier with an available small-molecule antagonist already in clinical development and with an established safety profile (ENB-003^66,67^; <u>NCT04205227</u> and BQ-788^68^; <u>NCT02442466</u>). Importantly, expression pharmacology confirmed increased EDNRB protein in stromal compartments of IMF-predominant patient biopsies as compared to QMF (**Fig. 4k; Extended Data 7)**, providing *in situ* validation. Together, these data position EDNRB as a high-confidence, clinically actionable target within disease-associated stromal signaling. In parallel, gene set enrichment analyses highlighted activation of JAK–STAT signaling in IMFs (**Fig. 1f**), providing a clinically relevant benchmark because the JAK inhibitor upadacitinib (UPA) is an FDA-approved therapy for moderate-to-severe, treatment-refractory CD and UC, often used as salvage therapy^69–77^.

We first compared EDNRB as a novel actionable node, and benchmarked readouts in human NAMs against UPA (see workflow, **Extended Data Fig. 8a**). EDNRB antagonism (using BQ-788) broadly suppressed IMF-associated secretory programs, reducing SASP cytokines and chemokines across multiplex assays (**Fig. 5a; Extended Data Fig. 8b**). This effect extended to the defining IMF chemokine CXCL14, which was significantly reduced at the protein level (**Fig. 5b**). While JAK inhibition attenuated select inflammatory mediators, its effects were partial and context-dependent, in contrast to the more uniform suppression achieved with EDNRB blockade. For example, while both inhibitors reduced CXCL14, IL1β and IL8 (**Fig. 5b**), only BQ-788, not UPA reduced levels of TNFα (**Extended Data Fig. 8b**).

**Figure 5.**
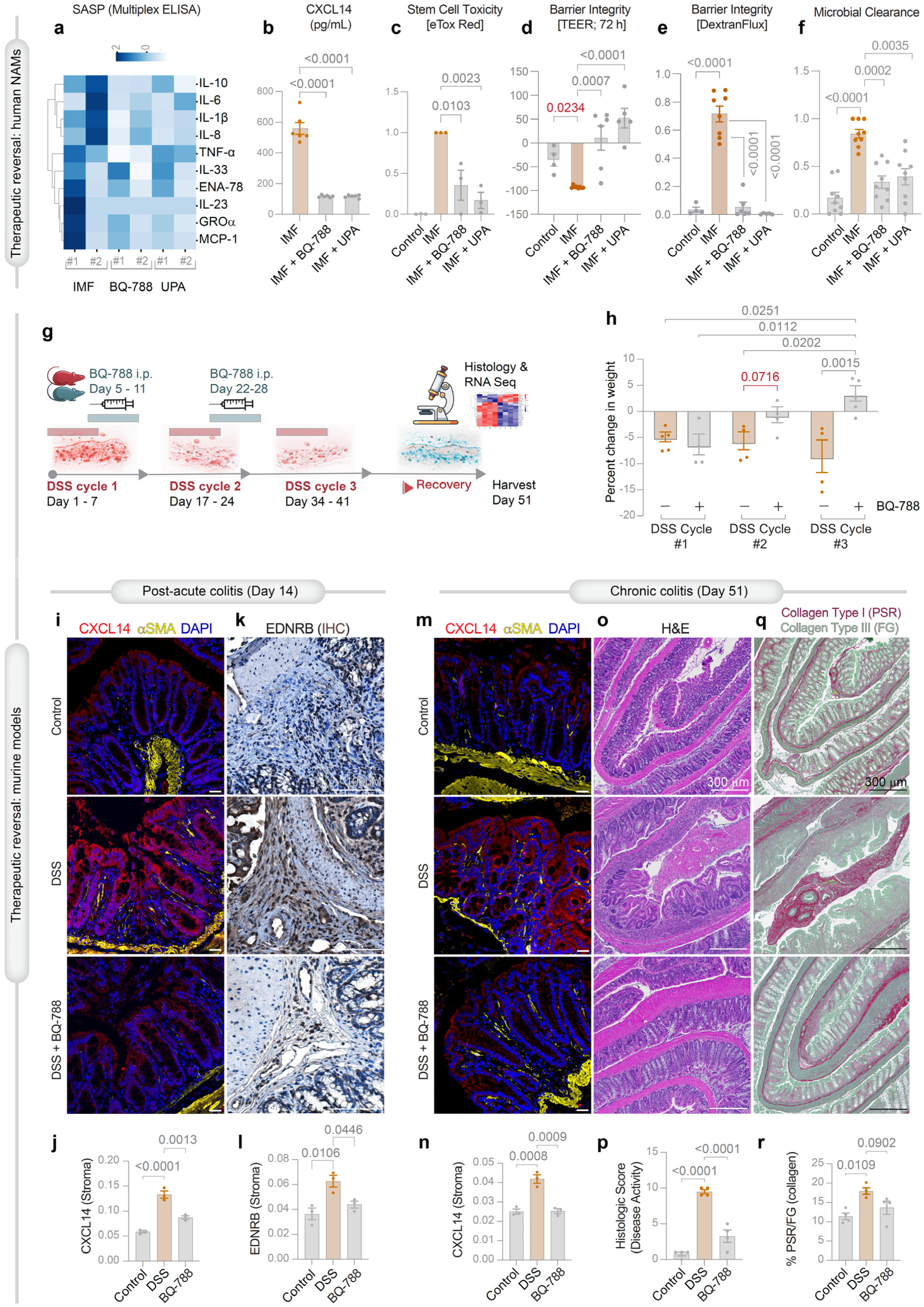
EDNRB antagonism restores stromal and tissue phenotypes. **a–f, Reversal of IMF-associated phenotypes in human NAMs** (see **Extended data 8a** for workflow). **a**, Multiplex electrochemiluminescence (MSD) profiling of SASP factors in conditioned media from IMFs treated with EDNRB antagonist (BQ-788) or JAK inhibitor (upadacitinib, UPA), shown as a heatmap of cytokine and chemokine secretion. See **Extended data 8b** for individual bar plots. **b**, CXCL14 secretion measured by ELISA. **c**, Organoid toxicity quantified by eTox Red incorporation. See **Extended data 8c** for representative images. **d**, Barrier integrity assessed by transepithelial electrical resistance (TEER) over 72 h. See **Extended data 8d** for time course measurements (0-72 h). **e**, FITC-dextran permeability across epithelial monolayers. **f**, Macrophage-mediated microbial clearance, quantified as surviving bacterial colony-forming units (CFUs). Results represent findings of conditioned media from independent donor-derived cultures; n = 2 unique IMF; 3-5 unique passages/subject. **g–h, *In vivo* therapeutic intervention in DSS colitis. g**, Experimental design of repeated DSS-induced colitis with EDNRB antagonist (BQ-788) administered intraperitoneally (i.p.) during defined windows across cycles. See also **Extended data 9** for rationalization of the timing of therapeutic initiation. **h**, Percent change in body weight across DSS cycles. Data represent independent biological replicates. **i–r, Tissue-level reversal of stromal activation and pathology. i, m**, Immunofluorescence staining of CXCL14 (IMF-associated; red), αSMA (stromal cells; yellow), and nuclei (DAPI; blue) in post-acute (day 14, **i**) and chronic (day 51, **m**) colitis. Scale bars, 20 µm. **j,n**, Quantification of CXCL14 signal intensity. **k-l**, EDNRB expression by immunohistochemistry. See **Extended Data Fig. 8b-c** for appropriate staining controls. **o-p**, Hematoxylin and eosin (H&E) staining showing tissue architecture and inflammation. **q-r**, Collagen deposition assessed by Picrosirius Red (PSR)/Fast Green (FG) staining. Data represent independent biological replicates. See also **Extended data 10**. *Statistics*: Bars denote mean ± S.E.M. Statistical comparisons were performed using one-way ANOVA (grey font) or two-sided, unpaired t-tests (red font), with exact *p* values shown in panels.

We next assessed functional rescue across epithelial and immune readouts. Conditioned media from BQ-788-treated IMFs markedly reduced epithelial toxicity, restoring organoid viability (**Fig. 5c; Extended Data Fig. 8c**). Barrier function was similarly rescued, with restoration of TEER and reduction of paracellular permeability (**Fig. 5d,e; Extended Data Fig. 8d**). In parallel, macrophage antimicrobial function improved, with enhanced clearance of bacteria following exposure to conditioned media from BQ-788-treated IMFs (**Fig. 5f**). Across these orthogonal assays, EDNRB antagonism with BQ-788 consistently reversed IMF-driven epithelial and immune dysfunction, comparable to those observed from JAK inhibition with UPA.

To test therapeutic relevance *in vivo*, we used a chronic Dextran Sulfate Sodium (DSS)-induced colitis model^78,79^ comprising repeated cycles of DSS exposure and recovery to recapitulate persistent inflammation, weight loss, and fibrosis seen in human IBD. Intervention timing (**Fig. 5g**) was guided by transcriptionally resolved stromal state dynamics (**Extended Data Fig. 9a**). IMF and QMF programs were suppressed during acute injury and emerged during remodeling phases, progressively accumulating with repeated cycles (**Extended Data Fig. 9b**) and paralleling induction of stromal drivers such as *CXCL14* and *ADAMDEC1* (**Extended Data Fig. 9c-d**). By contrast, *EDNRB* expression peaked during acute inflammation and declined during remodeling phases (**Extended Data Fig. 9e**), providing a rationale for temporally targeted antagonism. Accordingly, BQ-788 was administered during defined post-injury windows, mimicking treatment during clinical ‘flares’. This strategy attenuated disease severity, with improved weight trajectories and increased resilience to repeated injury (**Fig. 5g,h**).

At the tissue level, EDNRB antagonism reversed key features of stromal activation and pathology. CXCL14⁺ stromal compartments were markedly reduced in both post-acute and chronic phases of colitis (**Fig. 5i,j, m,n; Extended Data Fig. 10a-d**), consistent with suppression of IMF states. EDNRB induction in acute colitis and its suppression post-treated was also confirmed (**Fig. 5k,l**). Histologic analyses demonstrated restoration of epithelial architecture and reduction of inflammatory infiltrates (**Fig. 5o,p; Extended Data Fig. 10e**). Collagen deposition and fibrotic remodeling were similarly attenuated, as assessed by Picrosirius Red/Fast Green staining (**Fig. 5q,r; Extended Data Fig. 10f**).

Together, these data establish EDNRB as a tractable and high-confidence therapeutic target within the IMF program. Pharmacologic inhibition of EDNRB reverses stromal activation, restores epithelial barrier integrity, and rescues innate immune function across human NAMs and in mice. Thus, targeting a stromal state-defining node appears to achieve coordinated, multi-compartmental rescue, positioning EDNRB antagonism as a precision strategy to reprogram pathogenic stroma in IBD.

### An EDNRB response signature predicts therapeutic reprogramming from recalcitrance to remission

We next translated EDNRB-reversible IMF biology into a predictive framework for patient stratification. We derived a stromal state–specific EDNRB response signature by intersecting transcriptional changes induced by EDNRB antagonism in mice with human IMF programs (**Fig. 6a**). This yielded a refined 15-gene signature from 815 EDNRB-responsive genes (**Supplementary Data 4**; **Extended Data Fig. 11a,b**) and 224 IMF-defining genes (**Supplementary Data 2**), distilling a mechanistically anchored, target-linked biomarker of pathogenic stroma.

**Figure 6.**
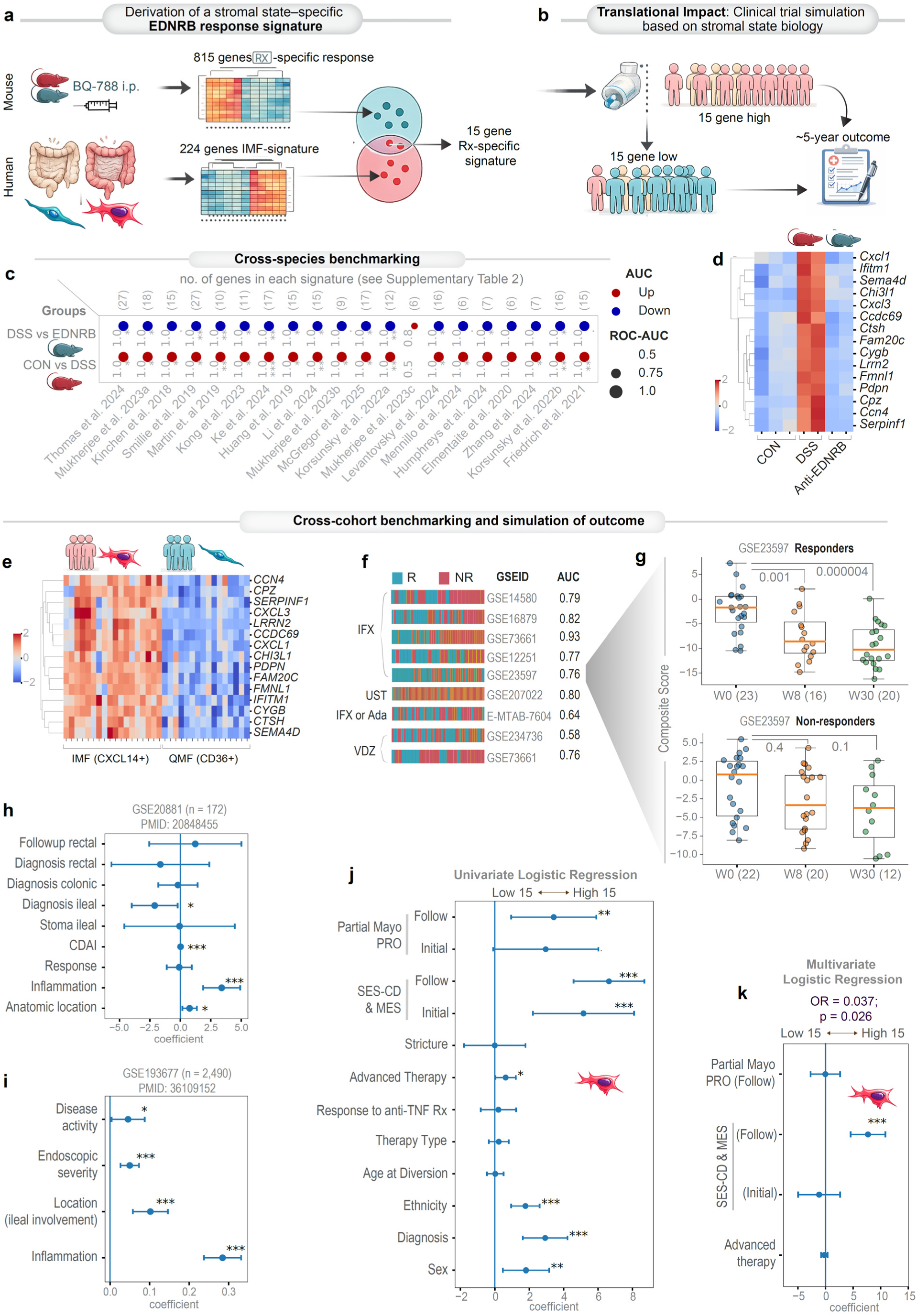
A stromal state–derived EDNRB response signature predicts therapeutic outcome. **a, Derivation of a stromal state–specific EDNRB response signature**. Gene expression profiling was performed following pharmacologic EDNRB antagonism (BQ-788) in murine models and integrated with human stromal transcriptional programs. Differential expression analysis identified an EDNRB response gene set (815 genes; see **Supplementary Data 4, Extended Data 11**) and an IMF state–specific program (224 genes; see **Supplementary Data 2**). Intersection of these datasets yielded a 15-gene EDNRB response signature enriched within the IMF stromal state. **b, Conceptual framework for clinical translation.** Patients are stratified based on expression of the stromal EDNRB response signature (high versus low 15-gene score), enabling simulation of predicted therapeutic outcomes over approximately five years based on stromal state biology. **c–d, Cross-model validation of EDNRB pathway activity. c**, Benchmarking of EDNRB-associated transcriptional responses across trajectory-defining stromal clusters from independent single-cell and bulk transcriptomic datasets. Dot size indicates ROC–AUC performance, and color denotes directionality of regulation. **d**, Heatmap of representative genes demonstrating modulation of inflammatory and stromal activation programs across control, DSS-induced colitis, and EDNRB antagonist–treated conditions. **e**, Heatmap showing enrichment of the 15-gene signature in CXCL14⁺ inflammatory myofibroblasts (IMFs) compared with CD36⁺ quiescent myofibroblasts (QMFs). **f–g, Prediction of therapeutic response across independent cohorts. f**, Application of the EDNRB response signature across publicly available transcriptomic datasets of inflammatory bowel disease treated with biologics (IFX, infliximab; UST, ustekinumab; VDZ, vedolizumab). Classification performance is summarized by ROC–AUC values. **g**. Longitudinal changes in composite signature scores among responders and non-responders over treatment timepoints (baseline, week 8, week 30) reflect differential dynamics of the stromal response program in responders vs. non-responders. See also **Extended Data 12a** for classification accuracy using the superset of IMF state–specific program (all 224 genes). **h–i, Association with clinical disease features.** Cross-sectional analyses across two independent cohorts linking the stromal EDNRB response signature with clinical disease features. Multivariate regression models were constructed using covariates identified in univariate analyses (see **Extended Data 12b-c**), demonstrating robust associations between the stromal response program and measures of disease activity and inflammation. **j–k, Prospective outcome modeling.** Univariate logistic regression analyses (j) linking stromal signature expression (low versus high 15-gene score; base variable) with patient characteristics and clinically relevant outcomes (see ***Methods***). Multivariate regression models (k) adjusting for clinical covariates demonstrate that a high 15-gene stromal signature at the time of tissue diversion predicts persistent worsening of endoscopic outcomes at follow-up (Simple Endoscopic Score for Crohn’s Disease [SES-CD] and Mayo Endoscopic Subscores [MES] for ulcerative colitis; see ***Methods***). Conversely, a treatment-associated low 15-gene signature identified patients protected from worsening disease (O.R = 0.037, 95% CI 0.002–0.88, Fisher’s exact P = 0.026). See **Supplementary Data 1** (’15 gene score’ column) for gene scores. PRO, patient reported outcomes. *Statistics*: Transcriptomic analyses integrate multiple independent cohorts and publicly available datasets. ROC–AUC values quantify classification performance. p-values were obtained using two-sided, unpaired Welch’s t-tests comparing score distribution between groups. Ordinary least squares models report coefficients with corresponding significance levels. Points represent regression coefficients with 95% confidence intervals; significance is indicated in panels. *P* values indicated in panels; P < 0.05 (*); <0.01 (**); <0.001 (***). Data visualization reflects normalized gene expression or composite signature scores as indicated.

We next established cross-species and cross-cohort robustness of EDNRB-reversible stromal states. Nearly all independently annotated stromal signatures were strongly induced by DSS, with consistent classification performance and preserved directionality (**Fig. 6c***-bottom*). EDNRB antagonism reversed these disease-associated programs toward a control-like state (**Fig. 6c***-top*), consistent with its suppressive impact on inflammatory and immune activation pathways (**Extended Data Fig. 11c**). Importantly, the 15-gene signature was selectively enriched in CXCL14⁺ IMFs relative to CD36⁺ QMFs (**Fig. 6e**), confirming its specificity for the pathogenic stromal compartment.

We then tested clinical utility across independent IBD cohorts treated with biologics (including infliximab, ustekinumab, vedolizumab). The 15-gene signature robustly predicted therapeutic response with high ROC–AUC across studies (**Fig. 6f**) and outperformed the full 224-gene IMF program in most instances (**Extended Data Fig. 12a**), indicating that mechanistic reduction enhances precision. Longitudinal analyses revealed that responders showed progressive normalization of the signature, whereas non-responders remained persistently elevated (**Fig. 6g**), reflecting failure to resolve stromal pathology. Cross-sectional analyses further linked higher signature scores to increased inflammation, disease activity, and endoscopic severity (multivariate analyses, **Fig. 6h,i**; univariate analysis, **Extended Data Fig. 12b-c**).

Finally, in our prospective cohort, high baseline expression of the 15-gene signature strongly predicted adverse outcomes. In univariate models, elevated signature scores tracked with persistent disease activity (**Fig. 6j**). In multivariate models adjusting for clinical covariates, the stromal signature emerged as a robust independent predictor of persistent endoscopic inflammation at follow-up. Compared with individuals in the highest quartile, those in the lowest quartile were significantly more likely to achieve mucosal healing. The odds of progressively worsening endoscopic scores were markedly reduced (O.R = 0.037, 95% CI 0.002–0.88, Fisher exact p = 0.026; **Fig. 6k**), indicating a strong protective association of the therapy-response signature (see ***Methods*** for clinical outcomes analysis).

Together, these data complete a translational arc from mechanism to precision medicine. A stromal, target-linked EDNRB response signature captures pathogenic IMF biology and predicts both therapeutic response and long-term disease trajectory, enabling precision stratification and nominating EDNRB-directed therapy to shift recalcitrant IBD toward remission.

## Discussion

IBD remains defined by an expansion of the therapeutic options without durable disease modification. Here, we identify stromal state plasticity as a central, targetable axis that links mechanism to outcome. This framework enables prediction, intervention, and reprogramming of disease trajectory. There are three major impacts of this study on the field.

First, we establish that human stromal states can be prospectively captured, propagated, and experimentally interrogated with high fidelity. By integrating biobanking with longitudinal outcomes, we move beyond static atlases to a system in which stromal programs are stable, reproducible, functionally testable and most importantly, determinants of disease-trajectory. Cross-cohort benchmarking using single-cell transcriptomic atlas derived digital signatures revealed that these states are conserved across cohorts, preserved in culture, and spatially encoded within tissue, providing a scalable human NAMs platform for the stromal compartment. Notably, the binary stromal organization (IMF vs QMF) parallels emerging insights from the epithelial compartment^34^, where patients also segregate into two dominant molecular subtypes in spite of their clinical and demographic heterogeneity. However, the axes are distinct: epithelial programs stratify fibrostenotic versus dysbiotic disease^34^, whereas stromal states segregate remitting versus recalcitrant trajectories. While both stromal states are observed across UC and CD, epithelial programs are largely restricted to CD^34^, suggesting that convergence of both epithelial and stromal drivers may underlie the most severe complications in IBD that are unique to recalcitrant CD, including transmural fibrosis, fistulizing disease and need for repeated surgeries. Consistent with this, we show that IMFs recapitulate transcriptional programs described in fistulizing CD^12^. Although this study does not resolve how epithelial and stromal subtypes interact, it addresses a central bottleneck in the field—how to move from single-cell description to causal and therapeutic insight—and positions human stroma as an experimentally tractable human NAM platforms for basic and translational discovery.

Second, our findings reframe stromal biology as a population-level equilibrium, in which the balance between inflammatory (IMF) and quiescent (QMF) myofibroblast states governs tissue fate. Although both states coexist within individual tissues and cultures, one predominates, and the dominant state dictates phenotype and clinical trajectory. IMFs emerge as senescent, metabolically sustained, and immunomodulatory effector population that couple epithelial injury with immune dysfunction, while QMFs represent a homeostatic population that maintains structural integrity and promotes repair. This “population game” mirrors principles observed across fibrotic diseases in multiple organs^19,24,80–82^, where emergence of conserved stromal activation states precedes and drives pathology. Within this framework, recalcitrant disease can be viewed as a non-healing wound state, marked by persistence and dominance of pathogenic IMFs that prevent restoration of tissue homeostasis. This state is sustained by a self-reinforcing loop of barrier failure and impaired microbial clearance. While not fully resolved, our data provides mechanistic clues into how such loops might be sustained or driven. SASP cytokines can directly disrupt epithelial junctions and permeability^83,84^; chemokines such as MCP1, together with stromal contractility^85^, may amplify immune cell recruitment beyond classical gradients. Conceptually, this shifts the therapeutic goal from eliminating cells to reprogramming stromal states, restoring balance within the compartment to convert a chronic non-healing wound into effective mucosal repair.

A key candidate linking stromal state to these downstream effects is CXCL14, a defining IMF chemokine that connects stromal activation to immune modulation, including recruitment of monocytes, NK cells and dendritic cells^86,87^. CXCL14 also functions as an autocrine factor for fibroblast growth, migration and multi-modal epithelium stimulatory activities^88^, and is expressed in the epithelium^89^ (consistent with our own observations in mice and patients), suggesting bidirectional stromal–epithelial crosstalk^90^. However, CXCL14 is unlikely to act alone. We posit that it may account for a subset of the paracrine effects we observe, with the broader phenotype arising from combinatorial signaling. For example, emerging evidence in hematopoietic^91^ and non-hematopoetic^92^ stem cells suggests that CXCL14 compromises stem cell fitness, decreasing regenerative potential and even increasing vulnerability to ferroptosis through metabolic stress pathways. In parallel, CXCL14-driven recruitment and polarization of macrophages toward tolerant states^93,94^, could impair microbial clearance, while coordinated engagement of recruited immune effectors^86^ may be required to fully manifest barrier dysfunction. Although its precise causal contribution remains to be defined, the consistent enrichment of CXCL14 in IMFs (here, and in most prior studies; see **Table 1**), together with its detectability in the secretome, positions it as a measurable biomarker of stromal state and therapeutic response, with the potential to translate gut stromal biology into a clinically accessible signal.

Third, we translate this biology into a precision therapeutic framework. We showed that JAK inhibition, prioritized guided by pathway enrichment, partially attenuates inflammatory signaling within IMFs, providing a mechanistic explanation for its efficacy in refractory disease. While originally developed to target cytokine signaling cascades in immune cells^95^, our human NAMs data demonstrate that the stromal compartment is a *bona fide* target of JAK inhibitors, albeit with incomplete reprogramming of the pathogenic state. These findings mechanistically substantiate and align with observations in rheumatologic diseases^84,96,97^ and position JAK inhibitors as cell-agnostic modulators of inflammatory cytokine signaling. By contrast, EDNRB emerges as an IMF state-defining, druggable node within the highest-confidence disease genes, enabling coordinated reversal of stromal activation, epithelial injury, and immune dysfunction. The derivation of an EDNRB response signature further operationalizes this biology into a predictive tool for patient stratification and outcome, directly linking target engagement to clinical trajectory. Our data also highlights timing as a critical determinant of efficacy. Consistent with fibrosis paradigms across organs, intervention during active inflammation (‘flares’) and early remodeling phases is required to intercept disease before stromal persistence becomes entrenched. Within this framework, recalcitrant IBD reflects failure of stromal resolution -- a non-healing wound -- rather than incomplete immune suppression, suggesting periodic, state-guided EDNRB-targeted intervention during flares (as an add-on therapy) may be both efficient and effective, in contrast to continuous treatment, and may represent a radically different approach from maintenance therapy..

Importantly, EDNRB inhibitors are already in clinical development with established safety profiles (ENB-003^66,67^; <u>NCT04205227</u> and BQ-788^68^; <u>NCT02442466</u>), enabling immediate translational potential. Consistent with oncology studies where EDNRB inhibition enhances checkpoint efficacy by dismantling immunotolerant niches^66,67^, our identification of elevated PD-L1 in IMFs suggests a parallel stromal checkpoint axis in IBD that can be therapeutically reprogrammed. Simulated outcomes based on stromal-state biology suggest that such a first-in-class stromal intervention, perhaps as a single agent, or in combination with others, could translate into a 96.4% lower odds of recalcitrant disease and shift patients toward remission. Together, these findings position EDNRB targeting as a precision strategy to reprogram pathogenic stroma and durably alter disease course in IBD.

### Study Limitations

This study has several limitations that frame the next steps. First, although the prospective cohort is deeply phenotyped with longitudinal follow-up, its size is modest, contributing to wide confidence intervals in outcome modeling. Validation across larger, multi-center and multi-ancestry cohorts is needed to confirm the generalizability of the stromal axis and the EDNRB response signature. Second, while primary stromal cultures preserve dominant state programs and enable mechanistic interrogation, they simplify the in vivo microenvironment; future studies incorporating multi-compartment systems, *in situ* perturbations and orthogonal multiomics studies will be required to resolve intercellular dependencies and deconstruct secretome complexity. Third, although the IMF↔QMF axis captures a dominant and clinically relevant dimension, it does not exhaust stromal heterogeneity, and longitudinal tracking of state dynamics during disease evolution remains to be established. Additional intermediate or context-specific states, as well as lineage relationships and plasticity dynamics, remain to be defined and tracked during disease evolution, particularly in larger or tissue-diverse cohorts. Finally, while EDNRB targeting is supported by human NAMs, in vivo models, and computational outcome simulation, clinical validation is required to establish efficacy, timing, and integration with existing therapies, particularly for state-guided intervention during disease flares.

## Conclusions

Together, this work establishes stromal state plasticity as a unifying axis linking cellular state to disease trajectory, therapeutic response, and intervention in IBD. By converting an elusive fibrotic cell type into a tractable human system, we bridge stromal atlases to clinical action. Myofibroblast states emerge as conserved, predictive, pathogenic, and reversible, thereby positioning the stroma as both a biomarker and a therapeutic target. This shifts the paradigm from reactive immune suppression to predictive, mechanism-based reprogramming of disease, with the potential to alter the natural history of recalcitrant IBD.

### Experimental Model and Subject Details

#### Human subjects

For generating healthy and IBD MFs, patients were undergoing colonoscopy as part of routine care for the management of their disease at the University of California, San Diego IBD-Center, following a research protocol compliant with the Human Research Protection Program (HRPP) and approved by the Institutional Review Board (Project ID# 1132632: PI Boland and Sandborn). Histologically normal healthy colon samples were collected from patients presenting for screening colonoscopy or undergoing the procedure to evaluate non-inflammatory symptoms, such as rectal bleeding. Each participant provided a signed informed consent to allow for the collection of colonic tissue biopsies for research purposes to generate a biobank of cell products. Isolation and biobanking of organoids from these colonic biopsies were carried out using an approved IRB (Project ID # 190105: PI Ghosh) that covers human subject research at the UC San Diego HUMANOID™ Center of Research Excellence (CoRE). For all deidentified human subjects, information including age, gender, and prior history of the disease, was extracted from the medical record in accordance with HIPAA regulations. The study design and the use of human study participants was conducted in accordance with the criteria set by the Declaration of Helsinki.

#### Clinical data collection and outcomes analysis

A comprehensive set of clinical parameters were collected at the time of tissue diversion and tracked prospectively for outcomes analysis (see **Supplementary Data 1** for the complete list of clinical parameters). These features included patient demographics (i.e. age, sex, and ethnicity), disease activity indices (i.e. Simple Endoscopic Score for Crohn’s Disease [SES-CD]^98^ or Mayo Endoscopic Subscore [MES]^99^ for ulcerative colitis; patient-reported outcomes [PRO]^100^ or partial Mayo scores^101^), as well as therapeutic history, including use of advanced therapies and treatment response. To enable cross-disease comparisons, analogous clinical measures were harmonized across endoscopic activity scores (SES-CD and MES) and symptom-based measures (PRO and partial Mayo scores) using min–max scaling.

#### Sex as a Biological Variable

Both male and female patients were included in this study. Analyses were conducted without sex-based exclusion.

## Method Details (Experimental and Computational)

### Experimental Methods

For a list of key reagents, consumables, software programs and equipment see **Table 3**.

### Isolation of myofibroblasts from colonic mucosal biopsies of healthy and IBD subjects

In brief, intestinal crypts were dissociated from tissues by digesting with collagenase type I (2 mg/mL solution containing gentamicin 50 µg/mL). The plate was incubated in a CO_2_ incubator at 37°C, mixing every 10 min with shear force pipetting in-between incubations, while monitoring the release of single epithelial units from tissue structures by light microscopy. After 80-90% of the epithelial cells were released from the tissue, the collagenase was deactivated with wash media (DMEM/F12 with Glutamax, 10% FBS) and the dissociated cells were passed through a 70 μm cell strainer. The remaining tissue caught in the filter was transferred to a T25 flask and supplemented with fibroblast growth media [MEM; L-glutamine (200 mM); MEM non-essential amino acids (1X); Antibiotic-Antimycotic (1X), sodium pyruvate (100 mM), heat-inactivated FBS (10 %)]^30,31^. Mesenchymal cells migrated from the tissue to the flask and cell colonies developed within 5-9 days post-digestion. The colonies were passaged to maintain 2D cultures and were characterized and biobanked. All experiments were conducted using passages 7 or less.

### Flow cytometric identification of primary colon myofibroblasts

Briefly, 2x10^5^ cells were passed through 70 µm filter and stained in FACS buffer (PBS, 5% FBS, 2 mM Sodium Azide) with CD45 Alexa fluor 488 and CD90 primary antibodies (see **Table 3**) for 30 minutes. Cells were fixed using the Cyto-Fast Fix Perm buffer set according to the manufacturer’s instructions. Cells were then stained in 1x cytofast perm wash with αSMA (1:50) for 30 minutes. Cells were washed and resuspended in FACS then acquired on Guava easyCyte flow cytometer. Data was analyzed using FlowJo software.

### Donor tissue benchmarking by immunohistochemical and immunofluorescent staining

Formalin-fixed, paraffin-embedded (FFPE) mucosal biopsy specimens from donor patients were used for immunohistochemistry (IHC) and immunofluorescent (IF) tissue staining. Tissue sections (4 μm thickness) mounted onto poly-L-lysine–coated glass slides were subject to deparaffinization in xylene and rehydration through graded ethanol. Heat-induced epitope retrieval was performed using Tris–EDTA buffer (pH 9.0), in a pressure cooker at 95°C for 20 minutes. For IHC, endogenous peroxidase activity was quenched with 3% hydrogen peroxide for 5 minutes. Sections were then blocked for 1 hour at room temperature with 2.5% goat or 2.5% horse serum to reduce nonspecific binding. Slides were then incubated with primary antibodies diluted in blocking buffer against CXCL14 (1:150), CD36 (1:150), αSMA (1:150), or EDNRB (1:200) (see **Table 3**). IHC antigen detection was performed using a streptavidin–biotin detection system with 3,3′-diaminobenzidine (DAB) as the chromogen and hematoxylin as a counterstain. Brightfield images were acquired using a BioTek Cytation 10, processed with Fiji/ImageJ software (NIH), and analyzed/quantified using Qupath (v0.5). For IF staining, sections were washed after primary antibody incubation, followed by incubation with Alexa Fluor-conjugated secondary antibodies (1:500) (see **Table 3**) and nuclear counterstaining with DAPI (1:1000) for 2 h at room temperature in the dark. Slides were mounted using ProLong™ mounting reagent, coverslipped, and sealed prior to imaging. Fluorescent images were acquired using a Stellaris 5 Confocal Microscope (Leica Microsystems), processed with Fiji/ImageJ software (NIH), and analyzed/quantified using Qupath (v0.5).

### Quantification of stromal MF subpopulations in donor tissue

Quantification of MF subpopulations within donor-derived mucosal FFPE sections was performed using QuPath (v0.5)^102^. Stromal regions were manually annotated to define regions of interest (ROIs) and exclude epithelial compartments. Cell detection was performed using StarDist-based^103^ nuclear segmentation to generate total nuclear counts within annotated stromal regions. Dual-marker classifiers were then trained to identify CXCL14⁺αSMA⁺ and CD36⁺αSMA⁺ double-positive MF populations using supervised object classification within QuPath. Following classifier training, automated batch analysis was applied across all images to identify MF subpopulations. The abundance of CXCL14⁺αSMA⁺ and CD36⁺αSMA⁺ cells was quantified relative to total nuclear counts within the stromal ROI, allowing normalization of MF subpopulations across samples. For quantification of single-marker intensity within tissue (i.e., CXCL14 by IF and EDNRB by IHC), pixel intensity for fluorescence or DAB signal was measured using QuPath within stromal-defined regions and normalized to total cell number within the ROI, as described above.

### Confocal immunofluorescence staining

MFs were seeded on 96-well plates at 5x10^3^ cells/well and cultured under standard conditions. MFs were fixed in 4% paraformaldehyde in PBS for 30 minutes at room temperature, quenched with 0.1 M glycine for 5 minutes, and then blocked and permeabilized with PBS containing 1% BSA and 0.1% Triton X-100 for 30 minutes at room temperature. Primary antibodies against CXCL14 (1:2000), CD36 (1:2000), COL1A1 (1:2000) (see **Table 3**) were diluted in blocking buffer and incubated overnight at 4 °C followed by Alexa Fluor-conjugated secondary antibodies (1:500), nuclear counterstaining with DAPI (1:1000), and F-actin staining with phalloidan (see **Table 3**) for 2 h at room temperature in the dark. Slides were mounted using ProLong™ mounting reagent, and cover slips were applied and sealed prior to imaging. Fluorescent images were acquired using a Biotek Cytation 10 at single-cell resolution. Quantification was performed by normalizing marker fluorescence intensity to individual cells using a DAPI-defined nuclear mask, enabling per-cell analysis and accounting for differences in cell number using Gen5 imaging software.

### Assessment of cellular senescence by SPIDER β-Galactosidase assay

MFs were seeded on 96-well plates at 5x10^3^ cells/well and cultured under standard conditions followed by incubation with the 100nM Bafilomycin A1 for 1 h in HBSS (without Ca2+/Mg2+) to inhibit endogenous β-galactosidase activity. MFs were washed using HBSS and incubated with 1 μM of SPIDER-βGal and 100nM Bafilomycin A1 (see **Table 3**) for 60 min at 37°C. Cells were resuspended in FACs buffer then filtered through 70 μm cell strainer and immediately acquired on a Guava easyCyte flow cytometer. Data was analyzed using FlowJo software. For imaging-based confirmation, SA-βGal signal was also assessed after seeding in 96-well plates and treated as described above with Bafilomycin A1 and SPIDER-βGal. Fluorescent images were acquired using a Biotek Cytation 10 at single-cell resolution. Quantification was performed by normalizing SA-βGal marker fluorescence intensity to individual cells using a DAPI-defined nuclear mask, enabling quantification on a per-cell basis using Gen5 imaging software.

### Assessment of DNA damage by flow cytometry

DNA double-strand breaks (DSBs) were measured by flow cytometry by detecting γ-H2AX in MFs after modifying the published methods^104^. Briefly, CD MFs were washed using FACS buffer (PBS, 5% FBS, 2 mM Sodium Azide) and fixed/stained using the Biolegend Cyto-Fast Fix Perm buffer set according to the manufacturer’s instructions. Cells were washed in Cyto-Fast Perm wash solution before incubation with the primary antibody anti-γ-H2AX (1:100) (see **Table 3**) for 30 min at room temperature. Cells were washed with Cyto-Fast Perm wash solution, followed by incubation with secondary antibody Alexa Fluor 488 conjugated goat anti-mouse IgG and propidium iodide for 30 min in the dark at room temperature. Cells were washed with Cyto-Fast Perm Wash Solution, resuspended in FACS buffer and data was acquired on a Guava easyCyte flow cytometer. Data was analyzed using the FlowJo software.

### Assessment of cell proliferation by Ki67 staining

MFs were grown for 72 hr in 8 chamber slides with a seeding density of 3.9x10^4^ cell/well followed by fixation, permeabilization as described above then incubated overnight with Ki67 (1:1000) primary antibody (see **Table 3**). Z stack images were acquired using confocal microscopy. From the acquired maximum intensity projection images, the number of Ki67 positive cells/total number of cells was used to generate the % of Ki67 positive cells. Cells were counted manually with 15 random fields of view, representative of the overall staining observed in each sample.

### Assessment of apoptosis by Annexin V

Apoptosis was assessed using eAnnexin V Red (see **Table 3**) and live-cell imaging on an RTCA eSight instrument (Agilent). MFs were seeded on 96-well plates at 5x10^3^ cells/well and cultured under standard conditions. At 48 h post-plating, eAnnexin V Red was added directly to the culture medium at a final concentration of 0.5 μg/mL according to the manufacturer’s instructions. Live-cell images were acquired using automated widefield imaging across all wells. Multiple fields per well were captured and analyzed using RTCA eSight analysis software to quantify Annexin V positive signal, which was normalized to total cell number per well.

### Assessment of cell bioenergetics by Seahorse

Cellular bioenergetics were assessed using a Seahorse XF Pro Analyzer (Agilent) to measure oxygen consumption rate (OCR) and extracellular acidification rate (ECAR). MFs were seeded in Seahorse XF96 cell culture microplates at 2x10^4^ cells/well and allowed to adhere overnight prior to analysis. On the day of the assay, culture media was replaced with Seahorse XF DMEM assay medium (see **Table 3**) supplemented with glucose, glutamine, and pyruvate, and cells were equilibrated in a non-CO₂ incubator for 1 h prior to measurement. Basal respiration and glycolytic activity were measured, followed by sequential injections of mitochondrial stress test compounds including oligomycin, FCCP, and rotenone/antimycin A according to the manufacturer’s protocol. OCR and ECAR were recorded in real time and analyzed using Wave software (Agilent). Bioenergetic parameters were normalized to total cell number per well using DAPI nuclear counterstain to allow comparison across conditions.

### Multiplex immunoassays for quantification of cytokines

Cytokines were quantified from conditioned media collected from MFs using customized Meso Scale Discovery (MSD) V-PLEX cytokine panels according to the manufacturer’s instructions. All data was obtained using a MESO QuickPlex SQ 120 instrument and analyzed using MSD® DISCOVERY WORKBENCH 4.0 software.

### Assessment of CXCL14 protein secretion by ELISA

CXCL14 protein levels were quantified by enzyme-linked immunosorbent assay (ELISA). Briefly, MFs were seeded on 96-well plates at 5x10^3^ cells/well and cultured under standard conditions. Conditioned media were collected, clarified by centrifugation to remove cellular debris, and analyzed using a commercially available CXCL14 ELISA kit (see **Table 3**) according to the manufacturer’s instructions.

### Assessment of PD-L1 expression

Measurement of surface PD-L1 expression on myofibroblasts was performed using flow cytometry. In brief, approximately 2x10^5^ MFs were washed in phosphate buffered saline (PBS), blocked using Human TruStain FcX (Fc Receptor) for 30 mins, followed by incubation with PerCP/Cy5.5-conjugated anti-human CD274 (see **Table 3**) for 1h on ice. After washing with 1% BSA in PBS, flow cytometry was performed using Agilent NovoCyte Quanteon with fluorescence-conjugated antibody and their controls followed. Data analysis was performed using FlowJo software and positive cells in percentage (%) were calculated.

### MF conditioned media (CM) production

5x10^5^ MFs were seeded in T25 flasks in 5 mL of fibroblast growth media (expanded above). Flasks were treated with 10 µM of EDNRB inhibitor (BQ-788), 10 µM of Upadacitinib (see **Table 3**) or left untreated 24 h after seeding. Treatments were replenished with concentrated stocks (10 µM final concentration) 24 h later. CM was collected 72 h post-seeding (48 h post-first treatment).

### Patient-derived organoid isolation and culture

Patient-derived organoids (PDOs) used in this study were isolated from colonic mucosal biopsy specimens obtained from healthy subjects using established crypt isolation protocols, as previously described^34^. In brief, intestinal crypts containing crypt-base columnar stem cells were enzymatically dissociated from colonic tissue using collagenase-based digestion, followed by mechanical disruption to release epithelial units. Digestion was quenched with wash medium, and epithelial fractions were filtered, centrifuged, and resuspended in Matrigel for three-dimensional culture. Matrigel domes were polymerized at 37 °C and overlaid with Intestigro^TM^ (see **Table 3**), a conditioned medium containing Wnt3a, R-spondin 3, and Noggin, supplemented with epidermal growth factor and small-molecule inhibitors to support epithelial stem cell survival and expansion, as previously reported^34^. Cultures were maintained at 37 °C with medium changes every 2-3 days and expanded or cryopreserved for biobanking.

### eTox stem cell cytotoxicity assay

Healthy PDOs were seeded on 96-well plates at 5x10^3^ PDOs/well and cultured in a 1:1 mixture of PDO growth medium and conditioned medium collected from MFs. Fibroblast growth media, used for routine MF culture, served as the control. PDO growth and morphology were monitored using live-cell imaging on a RTCA eSight instrument (Agilent) over the course of the experiment. After 96 hours, eTox Red cytotoxicity dye (see **Table 3**) was added to each well to a final concentration of 250 nM. Cytotoxicity was quantified by calculating the eTox Red fluorescence signal normalized to total PDO area per well, providing a measure of cell toxicity normalized to relative PDO number.

### Colonoid-derived monolayers and measurement of transepithelial electrical resistance (TEER)

Healthy EDMs were prepared by dissociating single cells from colonoids and plated in 96-well transwells with a 0.4 μm pore polyester membrane coated with diluted Matrigel (1:40) in 5% Tailor-2-Gro (see **Table 3**) as done before^34,105^. 24 h after incubation, media was completely replaced with MF CM or fibroblast growth media (control). High-throughput (HTP) assessment of TEER was carried out in an automated manner using the REMS AutoSampler (WPI) automated TEER measurement system. A WPI REMS-96C recording electrode was used to record TEER, which is compatible with a 96-well plate from Corning. The REMS-96C apical electrode was calibrated to measure TEER approximately 1 mm above the transwell membrane, and the transwell-read time was set to 5 sec/well. TEER was measured just before MF CM addition (0 h) and every 24 h following treatment for a total of 72 h. To mitigate TEER artifacts due to temperature fluctuations, the same read sequence is repeated every subsequent read. TEER recorded by REMS AutoSampler were saved as .txt files in raw TEER values (in Ωs).

### Assessment of barrier permeability of EDMs using FITC-dextran

After 72 h of treatment with MF CM, all media was replaced with HBSS and FITC-dextran (10 kD) (see **Table 3**) was added to the apical side at a 1:50 dilution. After 1 h of incubation with FITC-dextran, 50 µl of the basolateral supernatant was transferred to an opaque-black 96-well plate. Fluorescence was measured using excitation/emission 490 nm/520 nm with a Spark 20M Multimode Microplate Reader.

### Monocyte clearance assay

Adherent Invasive *Escherichia coli* strain LF82 (AIEC-LF82), isolated from the specimens of a Crohn’s disease patient, was obtained from Arlette Darfeuille-Michaud^61^. Bacterial clearance assay was performed as described previously^105^. Two days prior to infection, THP1 monocytes were seeded in THP1 media (Cytiva RPMI-1640+10%FBS+25nM PMA media) on a 24 well plate and incubated for 24 hours until successfully adhered and differentiated as macrophages. The following day, AIEC-LF82 was grown in LB broth for 6 hours under aerobic conditions and then for 14 hours under oxygen-limiting conditions overnight. THP1s were then cultured in a 1:1 mixture of THP1 growth medium and conditioned medium collected from MFs for approximately 16-18 hours prior to infection with AIEC-LF82 with a multiplicity of infection (MOI) of 30 for 1 hour. Gentamicin protection assay was performed after 1 hour infection with 200ug/mL of gentamicin for 90 minutes, followed by 3.5 hours of 50ug/mL of gentamicin. After 6 hours of clearance, THP1 cells were lysed in 1% Triton-x100 in PBS for 15 minutes, followed by serial dilutions in 1X PBS -1 through -5 and plated on LB Agar plates. Plates incubated for 16-18 hours and colonies were counted the next day to measure colony forming unit per ml (cfu/ml).

### Induction of acute and chronic colitis in mice

Male C57BL/6(B6) mice (7–8 weeks old) were purchased from Jackson Laboratories (Maine, USA) and acclimated for 2 weeks in pathogen-free conditions in the animal vivarium. To induce chronic fibrotic colitis, mice were given 2.5% DSS (molecular weight 36–50 kDa) orally in distilled water *ad libitum* for 7 days (from day 0 to 7). On day 7, they were switched to regular drinking water for 10 days, this cycle was repeated three times. Normal control mice received regular drinking water throughout the duration of the experiment. For induction of acute colitis, mice were orally administered with 2.5% DSS orally in distilled water *ad libitum* for 7 days (from day 0 to 7), followed by a 7-day recovery period. Mice were divided into three groups namely healthy (receiving normal drinking water), DSS only group and EDNRB-inhibitor (BQ-788) treated group (DSS administered and receiving 10 mg/kg dose of BQ-788, intraperitoneally, starting from days 5-11 of each of the three cycles).

### Evaluation of colitis

Body weight loss (scores: 0, none; 1, 0–5%; 2, 6–10%; 3, 11–15%; 4, 16–20%; 5, 21–25%; 6, 26–30%), stool consistency (scores: 0, normal stools; 1, soft stools; 2, liquid stools), hemoccult positivity and the presence of gross blood (scores: 0, negative fecal occult blood; 1, positive fecal occult blood; 2, visible rectal bleeding) were assessed daily for each mouse. The disease activity index (DAI) was calculated as a sum of the scores of the three parameters according to the scoring criteria as described in previous studies^106–109^.

### Evaluation of histology

The entire colon from each group (4 mice per group) was collected, and its length was measured immediately. The tissue was then fixed in 4% formalin, embedded in paraffin, and cut into thin cross-sections (4 µm thick). These sections were stained with hematoxylin and eosin (H&E) to assess inflammation. All stained sections were examined under a light microscope. For analysis, we focused on regions of the distal colon and evaluated and scored following signs of inflammatory cell infiltrate, epithelial changes and alteration to the mucosal architecture, as described previously^110^. The scores were quantified and statistically analyzed using GraphPad. To evaluate fibrosis and collagen deposition, additional sections were stained using Pico Sirius Red/Fast Green (PSR/FG)^111^, where Pico Sirius red stained the collagen deposition pink and fast green demarcated the non-collagen components as green. The difference in collagen deposition across treatment groups was quantified as Pico Sirius red positive stained area using ImageJ and analyzed using GraphPad,

### RNA isolation from mouse tissue

Mouse colonic tissue was harvested and immediately processed for RNA extraction. Tissue samples were first mechanically homogenized using a Precellys tissue homogenization system in TRI-Reagent, followed by RNA isolation using the Zymo Research Direct-zol RNA Miniprep Kit (see **Table 3**) according to the manufacturer’s instructions (see **Table 3**). RNA concentration and quality were assessed prior to downstream applications.

### Software and statistical analysis

All images were processed using ImageJ, QuPath, Gen5, or LAS X software, and assembled into Figure panels with Illustrator (Adobe Creative Cloud). All graphs were generated using GraphPad Prism v9.3.1. All experiments were repeated to confirm reproducibility (and biological and technical replicates are disclosed in each panel), and results were presented as average ± S.E.M. Methodologies for assessing statistical significance are disclosed in each figure panel and actual *p* values are displayed.

### Computational Methods

#### RNA sequencing and data processing

RNA sequencing libraries were generated using the Illumina TruSeq Stranded Total RNA Library Prep Gold with TruSeq Unique Dual Indexes (Illumina, San Diego, CA). Samples were processed following manufacturer’s instructions, except modifying RNA shear time to five minutes. Resulting libraries were multiplexed and sequenced with 100 basepair (bp) Paired End (PE100) to a depth of approximately 40 million reads per sample on an Illumina NovaSeq 6000. Samples were demultiplexed using bcl2fastq v2.20 Conversion Software (Illumina, San Diego, CA). Raw FASTQ files were trimmed and filtered using Trimmomatic to remove low-quality bases and adapter sequences. Reads were aligned to the human reference genome (GRCh38, Ensembl release 94) using STAR (v2.6.0a) with transcriptome alignment enabled. Transcript-aligned reads were quantified using RSEM (v1.3.3^112^) with the “–forward-prob 0” option. Gene-level transcripts per kilobase million (TPM) values from RSEM gene result tables were used for downstream analyses. We used TPM as a normalized RNA-seq quantification unit because it allows for accurate comparisons of gene expression across samples, as the sum of TPM values is identical in every sample; it enables identifying stable reference genes by evaluating low variability (low coefficient of variation) across experimental conditions^112,113^. Expression values were transformed using log2(TPM) for TPM > 1 and TPM – 1 for TPM ≤ 1. This transformation preserves dynamic range for highly expressed genes while minimizing inflation of low-abundance transcripts. Publicly available transcriptomic datasets were obtained from the Gene Expression Omnibus (GEO) repository and processed using the same normalization and transformation procedures to ensure analytical consistency across cohorts.

#### Principal component analysis (PCA)

Unsupervised principal component analysis (PCA) was performed, as done previously^114^, on log-transformed, normalized gene expression matrices to reduce dimensionality and evaluate global transcriptional structure. Prior to analysis, genes with low variance across samples were excluded to minimize noise. PCA was computed using singular value decomposition, and principal components were derived from the covariance matrix of scaled expression values. Samples were visualized in principal component space using the first two components. All analyses were conducted in Python using the scikit-learn package.

#### Differential expression analysis

Differential gene expression analysis was performed using raw count data processed through DESeq2^115^. Genes with an absolute log2 fold change ≥ 0.5 and an adjusted p-value ≤ 0.05 were considered differentially expressed genes (DEGs). Pathway enrichment analysis of DEG lists were conducted using the ShinyGO ^116^ database and its integrated algorithms.

#### Gene Set Enrichment Analysis (GSEA)

To contextualize transcriptional divergence between inflammatory (IMF) and quiescent (QMF) myofibroblast states, we performed preranked Gene Set Enrichment Analysis (GSEA)^117^ using gene-level t-statistics derived from IMF versus QMF comparisons. Enrichment was evaluated against MsigDB Hallmark and Reactome collections using 1,000 phenotype permutations. This approach enabled unbiased assessment of coordinated pathway-level shifts without imposing arbitrary fold-change thresholds. Normalized enrichment scores (NES), nominal p-values, and false discovery rates (FDR) were used to determine statistical significance.

#### StepMiner algorithm

*StepMiner* is an algorithm designed to detect stepwise transitions in time-series gene expression data^118^. It fits step functions to expression profiles by identifying the sharpest changes in signal, corresponding to gene expression switching events. The algorithm evaluates all possible step positions and calculates the average expression on either side of each step to define constant segments. An adaptive regression approach is then used to select the step position that minimizes the sum of squared errors between the observed and fitted data. The selected step is used as the *StepMiner* threshold. This threshold is used to convert gene expression values into Boolean values (0 or 1). A noise margin of 2-fold change is applied around the threshold to determine intermediate values, and these values are ignored during Boolean analysis. Finally, a regression test statistic is computed to assess the significance of the identified step transition as follows:

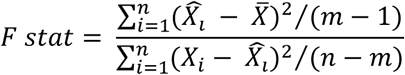

Where *X_i_* for *i* = 1 to *n* are the values, *X̂_l_* for *i* = 1 to *n* are fitted values. M is the degrees of freedom used for the adaptive regression analysis. *X̄* is the average of all the values: 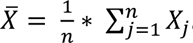. For a step position at k, the fitted values *X̂_l_* are computed by using 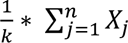 for *i* = 1 to *k* and 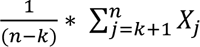 for *i* = *k* + 1 to *n*.

#### Composite gene signature analysis using Boolean network explorer

Boolean network explorer (BoNE)^119^ provides an integrated platform for the construction, visualization and querying of a gene expression signature underlying a disease or a biological process in three steps: **(i)** *<u>Binarization</u>*: The expression levels of all genes in these datasets are converted to binary values (high or low) using the *StepMiner* algorithm^118^. **(ii)** *<u>Normalization</u>*: Gene expression values are normalized using a modified Z-score approach centered on the *StepMiner* threshold [formula = (expr – SThr)/(3*stddev)]. **(iii)** *<u>Scoring</u>*: Normalized expression values for each gene in a given signature, are summed to generate a composite score, for the gene signature, representing the overall activity of the biological pathway in a sample.

As a modified Z-score, the composite score of a gene signature ranges from negative to positive values, reflecting the dynamic range of each gene in the signature. These composite scores enable the comparison of biological states between control and query groups within a dataset. However, comparisons between different gene signatures or across datasets are not valid, due to the lack of cross-signature normalization and dataset-specific normalization processes.

Samples are ranked by composite signature scores, and statistical differences between groups were assessed using Welch’s t-test (unpaired, unequal variances and sample sizes). Visualizations, including violin, swarm, and bubble plots were generated using the Seaborn Python package (version 0.10.1). Composite gene scores were computed using the COMPASS framework^120^ (https://compass.precsn.com/), which applies *StepMiner*-based thresholding and normalized aggregation of gene expression to derive sample-level pathway activity scores.

#### Measurement of classification strength or prediction accuracy

To evaluate classification strength and prediction accuracy, Receiver Operating Characteristic (ROC) curves were generated for each gene, along the lines we did previously^121,122^. These curves assess the performance of a binary classifier, e.g., high vs. low *StepMiner* normalized gene expression levels, across varying discrimination thresholds. ROC curves plot the True Positive Rate against the False Positive Rate at multiple threshold levels. The Area Under the Curve (AUC) quantifies the classifier’s ability to correctly distinguish between the various groups of samples. ROC-AUC values were computed using the Python Scikit-learn package. AUC values were visualized using bar or dot plots, with statistical significance assessed using Fisher’s exact test.

#### Expression heatmap

Z-normalized expression values for selected genes were visualized as a heatmap across all samples within the dataset using the Seaborn package (version 0.10.1).

#### Univariate and multivariate linear regression analysis

To quantify associations between transcriptional signatures and clinical or phenotypic variables, univariate and multivariate linear regression analyses were performed, exactly as done previously^35,121,123^, using ordinary least squares (OLS) models implemented in the Python statsmodels package. In univariate analyses, individual predictors were evaluated independently to assess their relationship with the outcome variable. Multivariate models incorporated relevant covariates, including demographic and clinical parameters, to control for potential confounding effects and to estimate independent contributions of each predictor. Clinical outcomes, including disease activity indices (patient-reported outcomes [PRO]^100^ or partial Mayo scores^101^) and endoscopic severity scores (Simple Endoscopic Score for Crohn’s Disease [SES-CD]^98^ or Mayo Endoscopic Subscore [MES]^99^ for ulcerative colitis), were first harmonized across scoring systems using min–max normalization to enable comparability across scoring systems and subsequently dichotomized into favorable and unfavorable states using consistent thresholds. Model assumptions, including normality and homoscedasticity of residuals, were evaluated using diagnostic plots. Regression coefficients, standard errors, and two-sided p-values were reported, with statistical significance determined at p < 0.05.

#### Odds ratio (OR) analysis

Associations between molecular features and clinical outcomes were evaluated using odds ratios (ORs) derived from 2×2 contingency tables. Clinical and demographic variables were dichotomized using predefined strategies. Stromal phenotype was analyzed as a binary variable (IMF vs QMF), whereas continuous gene scores for anti-EDNRB therapy were stratified using extreme quartiles (upper >75th percentile vs lower <25th percentile).

For each comparison, ORs were calculated from contingency table cross-products. Statistical significance was assessed using two-tailed Fisher’s exact test, and 95% confidence intervals (CIs) were estimated using the log-transformed standard error. To assess the protective impact of EDNRB, inverse OR weighting was applied using the inverse of the estimated odds ratio function relating exposure to the mediator, as previously described^124^. To reduce instability arising from sparse cell counts, particularly when a single observation can disproportionately influence estimates in small samples^125^, Haldane–Anscombe correction was applied when deemed appropriate.

## Supporting information

SOM

## Data availability

All materials are available from the lead contact with a completed Materials Transfer Agreement and patented technology agreement following the guidelines of the University of California, San Diego. The study protocol and statistical plan are available within the article and **Supplementary Information**. The raw data for preclinical experiments are available in **Source Data**, whereas the individual data from patients with each molecular subtype is available upon reasonable request from academic or qualified researchers affiliated with recognized institutions, strictly for the purpose of conducting ethically approved research. Applicants are required to submit a detailed research proposal, and declaration of non-conflict of interest to UC San Diego HUMANOID™ Center. All approved requestors will be required to sign a data access agreement that restricts data use solely to the approved research project and prohibits any further distribution. All transcriptomic datasets generated in this work have been deposited at NCBI GEO [GSE327844 and GSE328066]. Source data are provided with this paper. Any additional information required for reanalyzing the data reported in this work paper is available from Lead Contact upon request.

## Code availability

All code and computational workflows necessary to reproduce the analyses presented in this study are publicly available at: https://github.com/sinha7290/Myofibroblast.

## ACKNOWLEDGEMENTS

This work was supported by the Leona M. and Harry B. Helmsley Charitable Trust (to PG and BB). Other sources of support include-- National Institutes for Health (NIH) grants AI141630, UG3TR003355, UG3TR002968 and R01-AI155696 (to P.G). B.S.B. was supported by NIH K23DK123406 and P30DK120515. H.M.P. was supported by The California Institute of Regenerative Medicine Training Grant for Clinical Fellows through Sanford-Burnham Prebys-UC San Diego and a training grant from the NIH/NIDDK (T32 DK142622). S.S. was supported by the American Association of Immunologists Intersect Fellowship Program for Computational Scientists and Immunologists. GDK was supported through The American Association of Immunologists Intersect Fellowship Program for Computational Scientists and Immunologists and the American Heart Association (24CDA1268506). The authors acknowledge instrumentation resources at the UC San Diego Agilent Center of Excellence in Cellular Intelligence. The content is solely the responsibility of the authors and does not necessarily represent the official views of the Helmsley Charitable Trust or the National Institutes of Health. This publication includes data generated at the UC San Diego IGM Genomics Center utilizing an Illumina NovaSeq 6000 that was purchased with funding from a National Institutes of Health SIG grant (#S10 OD026929).

## AUTHOR CONTRIBUTIONS

C.T., H.M.P., S.S., M.M. and P.G. conceptualized the project. B.S.B and P.G supervised, and P.G acquired funding to support it. K.Z., H.N.L., J.N., W.S.J and B.S.B provided key resources for human subjects and were responsible for the selection and enrollment of patients into this study. CT, HMP, K C-P, MH, SSC, MM, C-CH, EM and G.D.K contributed experimentation and formal analysis. C.T generated the ‘living’ biobank. K C-P conducted all macrophage studies. S.S. performed all computational analyses. M.M., with assistance from EM and C-CH conducted all animal studies. H.M.P., S.S., and P.G. prepared figures for data visualization. P.G. wrote the original draft. H.M.P, S.S., C.T., B.S.B, and P.G. reviewed and edited the draft. All co-authors approved the final version of the manuscript.

---

**Extended Data Figure 1.**
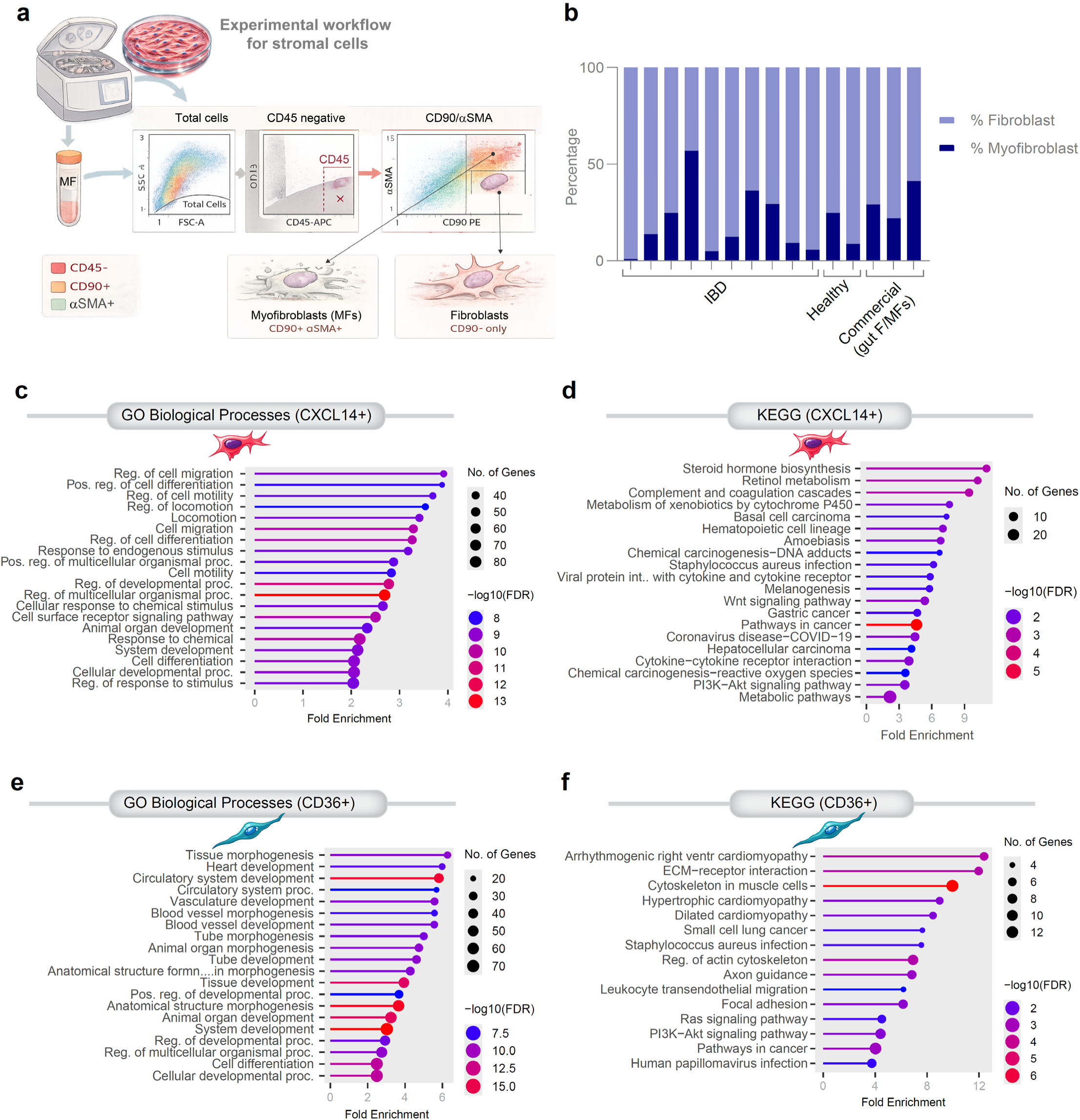
Isolation and characterization of stromal cell populations and pathway enrichment analyses. **a**, Experimental workflow for stromal cell isolation and identification. Tissue-derived single-cell suspensions were analyzed by flow cytometry. Cells were first gated on forward and side scatter to identify total cells, followed by exclusion of hematopoietic cells using CD45-negative gating. Stromal populations were subsequently resolved based on CD90 and αSMA expression, identifying myofibroblasts (MFs; CD90⁺ αSMA⁺) and fibroblasts (CD90⁺ αSMA⁻). **b**, Quantification of fibroblast and myofibroblast fractions across individual IBD samples and healthy controls enrolled into the current study, alongside commercially obtained positive controls (Lonza; Cat# CC-2902). Stacked bar plots show the relative percentage of fibroblasts and myofibroblasts within the CD45⁻ stromal compartment. **c–d**, Gene ontology (GO) and KEGG pathway enrichment analyses for genes upregulated in **CXCL14⁺ inflammatory stromal cells**. Enriched GO biological processes include cell migration, regulation of cell motility, response to endogenous stimulus, and cellular developmental processes (**c**). KEGG pathways enriched in the CXCL14⁺ group include complement and coagulation cascades, xenobiotic metabolism, cytokine signaling pathways, and PI3K–Akt signaling (**d**). **e–f**, Functional enrichment analyses for genes upregulated in **CD36⁺ quiescent stromal cells**. GO biological processes highlight tissue morphogenesis, vascular development, and circulatory system development (**e**). KEGG pathway analysis identifies enrichment of ECM–receptor interaction, cytoskeletal organization, focal adhesion, and Ras/PI3K signaling pathways (**f**). Bubble size indicates the number of genes contributing to each pathway and color denotes −log10(FDR).

**Extended Data Figure 2.**
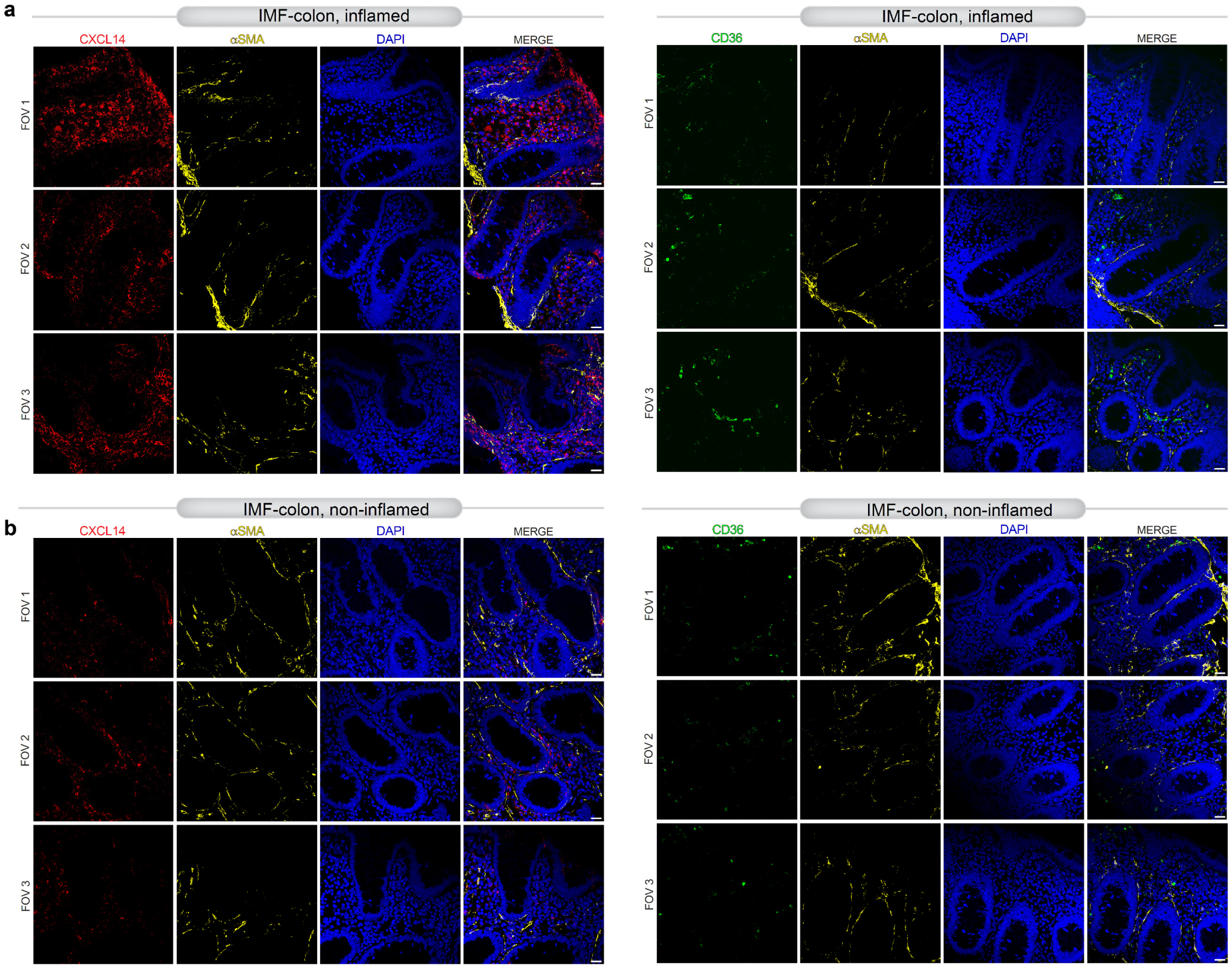
Donor tissue validation of IMF and QMF stromal states across inflammatory contexts in IMF-predominant patients. Representative immunofluorescence images of inflamed (**a**) and non-inflamed (**b**) colonic regions from IMF-predominant tissues stained for CXCL14 (IMF-associated; red) or CD36 (QMF-associated; green), together with αSMA (stromal cells; yellow) and nuclei (DAPI; blue). Multiple fields of view (FOV1–3) are shown per condition to demonstrate spatial consistency. Inflamed regions exhibit expansion of CXCL14⁺αSMA⁺ stromal compartments with relative paucity of CD36⁺ cells, whereas non-inflamed regions show reduced CXCL14 signals. Across all panels, merged images illustrate co-localization of stromal markers with αSMA⁺ cells. Representative images are shown from independent donor tissues with multiple fields of view per region and condition. Scale bars, 20 μm.

**Extended Data Figure 3.**
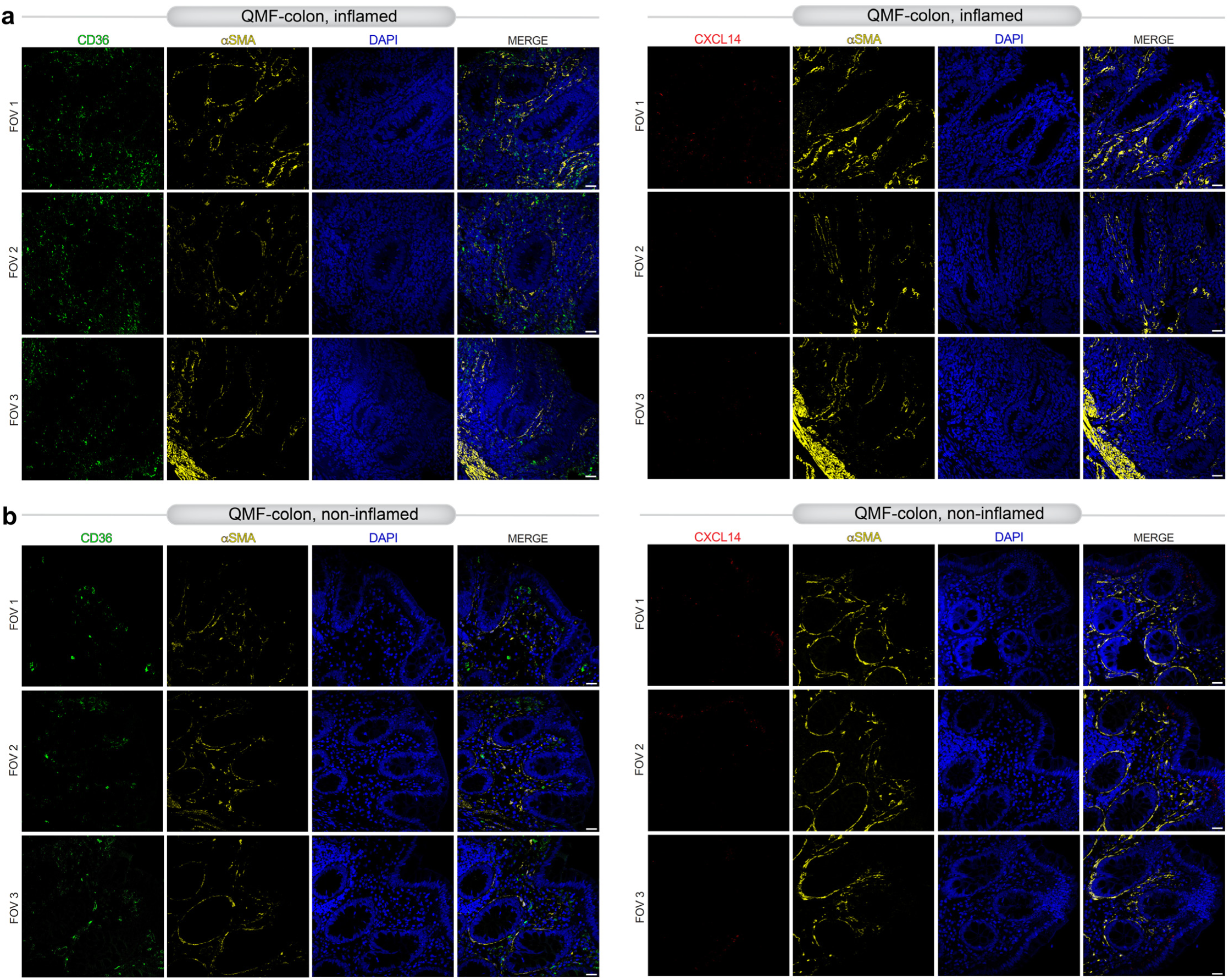
Donor tissue validation of IMF and QMF stromal states across inflammatory contexts in QMF-predominant patients. Representative immunofluorescence images of inflamed (**a**) and non-inflamed (**b**) colonic regions from QMF-predominant tissues stained for CXCL14 (IMF-associated; red) or CD36 (QMF-associated; green), together with αSMA (stromal cells; yellow) and nuclei (DAPI; blue). Spatial localization highlights compartmentalization of stromal states within the lamina propria and their relationship to epithelial crypt architecture. Across all panels, merged images illustrate co-localization of stromal markers with αSMA⁺ cells. Representative images are shown from independent donor tissues with multiple fields of view per region and condition. Scale bars, 20 μm.

**Extended Data Figure 4.**
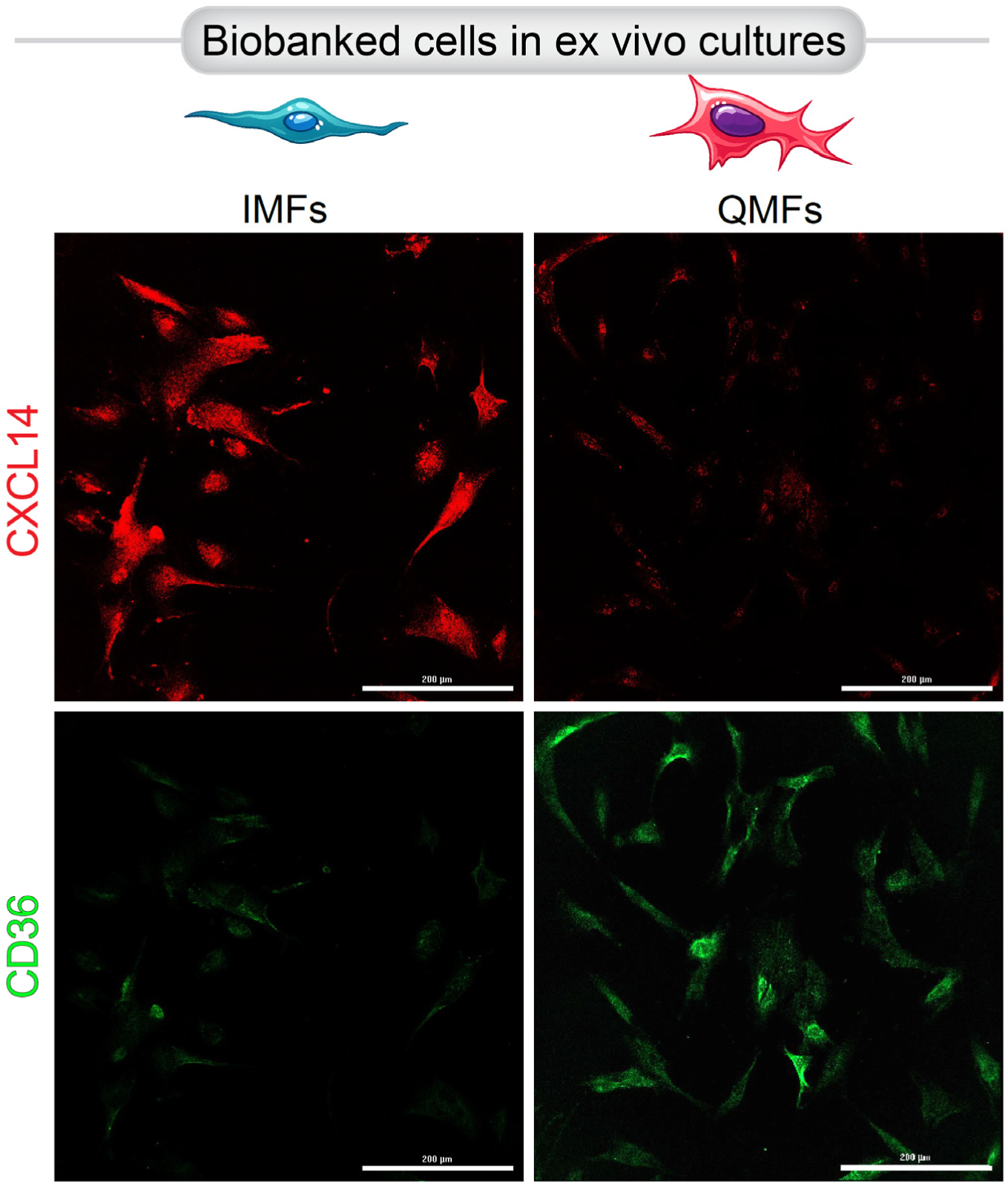
Donor-tissue benchmarking of IMF and QMF stromal states in culture. Representative immunofluorescence images of cultured, biobanked primary IMF or QMF MFs stained for CXCL14 (red) or CD36 (green). Scale bar, 200 µm. Quantification is shown in Fig 2f.

**Extended Data Figure 5.**
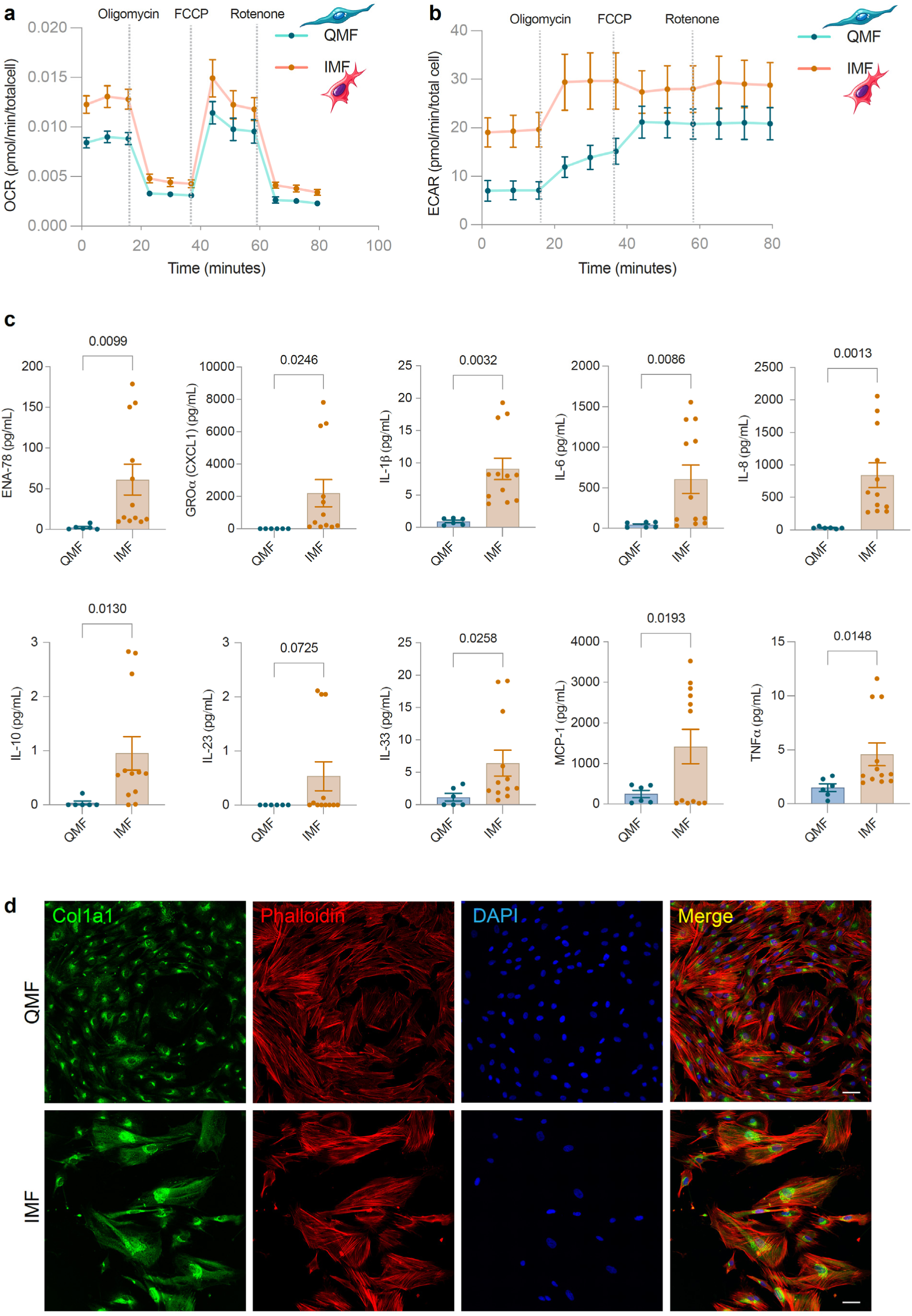
Metabolic and extracellular matrix features of QMF and IMF stromal states. **a–b, Real-time metabolic flux analysis**. Seahorse extracellular flux analysis compares quiescent myofibroblasts (QMFs) and inflammatory myofibroblasts (IMFs). Oxygen consumption rate (OCR, **a**) over time during sequential metabolic perturbations, reflecting mitochondrial respiration. IMFs exhibit consistently elevated OCR relative to QMFs across the assay, consistent with increased oxidative metabolism. FCCP denotes Carbonyl cyanide-4-(trifluoromethoxy)phenylhydrazone, a potent mitochondrial uncoupler. Extracellular acidification rate (ECAR, **b**) measured over time, reflecting glycolytic activity. IMFs display increased glycolytic flux compared with QMFs, indicating an energetically active metabolic state. **c**, Multiplex electrochemiluminescence immunoassay (MSD platform; Figure 3g) used to quantify SASP-related inflammatory mediators in stromal cultures. Each point represents an independent donor-derived culture; n = 4 unique IMF and 2 unique QMF subjects; 3 unique passages/subject. **d**, Representative immunofluorescence images of QMFs and IMFs stained for collagen type I (Col1a1; green), F-actin cytoskeleton (phalloidin; red), and nuclei (DAPI; blue). Merged images illustrate enhanced collagen deposition and altered cytoskeletal organization in IMFs relative to QMFs, consistent with an activated, matrix-producing stromal phenotype. Scale bar, 25 µm. Data in **a–b** represent mean ± S.E.M. from independent donor-derived cultures measured across technical assay replicates. Statistical comparisons were performed using two-sided, unpaired t tests; exact *P* values are shown. Representative images in **d** are shown from multiple biological replicates with multiple fields of view per condition.

**Extended Data Figure 6.**
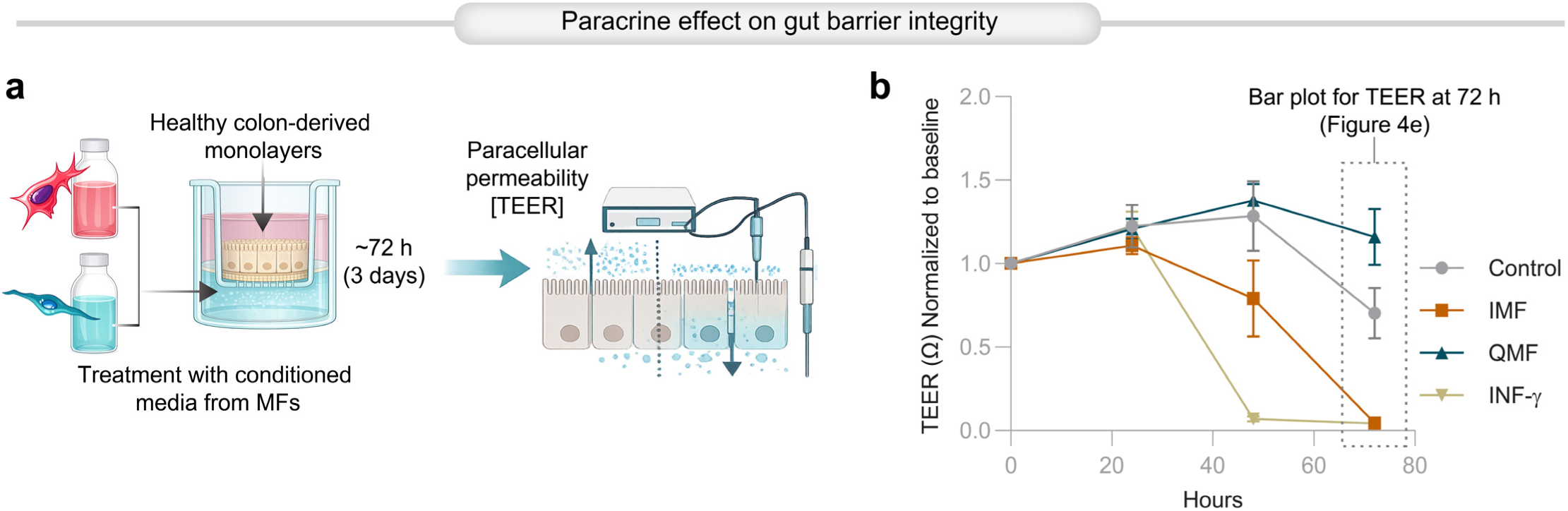
Cross-tissue validation of myofibroblast states and extended analysis of epithelial barrier dysfunction. **a–b**, Extended analysis of epithelial barrier function corresponding to Fig. 4d**–f**. Schematic of workflow (**a**) for measurement of paracellular permeability by transepithelial electrical resistance (TEER). Time-course (**b**) of TEER (0–72 h) normalized to baseline across control, QMF, IMF, and IFNγ-treated conditions, demonstrating progressive barrier loss induced by IMF-conditioned media and preservation by QMF-conditioned media. These data validate that IMF-associated programs are conserved across tissues and mechanistically linked to epithelial barrier disruption. *Statistics*: Line plots show mean ± S.E.M.

**Extended Data Figure 7.**
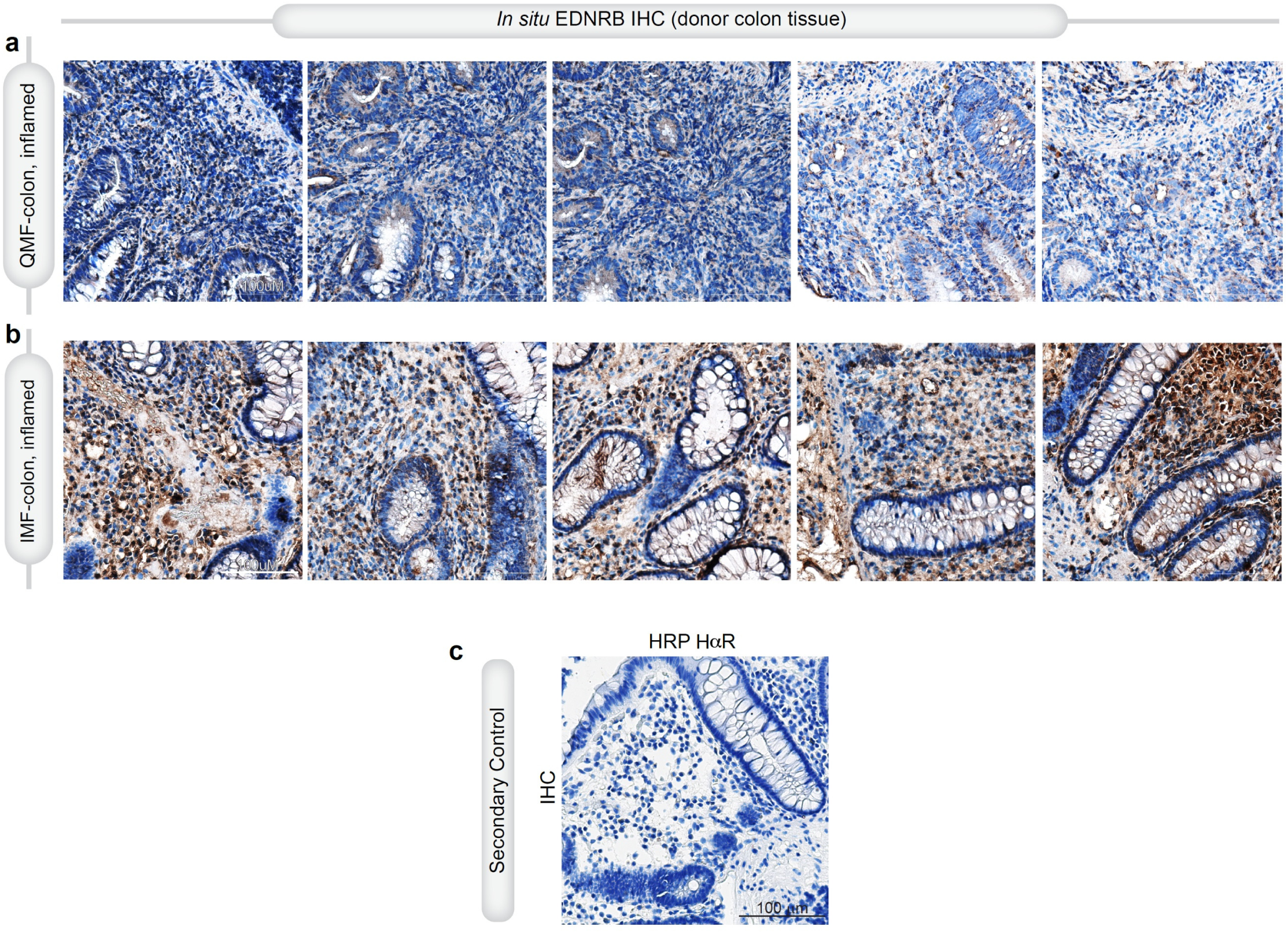
Stromal EDNRB expression is upregulated in IMF colonic mucosal tissue sections. **a-b**, Representative immunohistochemistry (IHC) images and quantification of inflamed colonic regions from QMF (**a**)- or IMF (**b**)-predominant tissues stained for EDNRB. Scale bar 100 µm. **c**, IHC negative control (HRP-conjugated secondary antibody only) confirming specificity of staining and absence of nonspecific signal. Scale bar 100 μm. *Reproducibility*: Representative images are shown from independent patients with multiple fields of view per condition. All staining and imaging were performed using identical acquisition and processing parameters across groups.

**Extended Data Figure 8.**
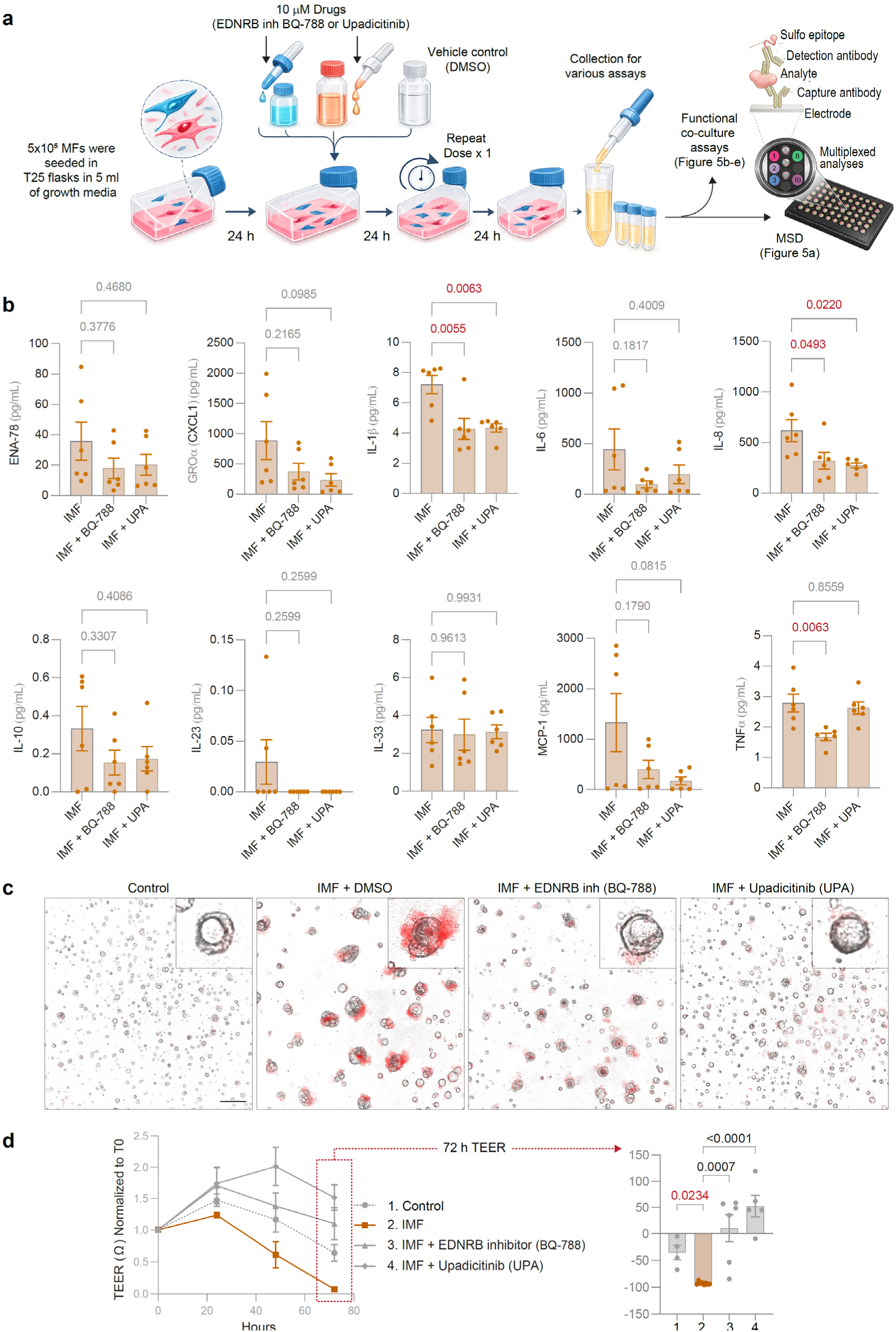
Pharmacologic modulation of IMF-derived paracrine programs in human NAMs. **a**, Workflow for generation of conditioned media from primary human IMFs. Cells were seeded in T25 flasks, treated with EDNRB antagonist (BQ-788), JAK inhibitor (upadacitinib, UPA), or vehicle (DMSO), with repeat dosing at 48 h, followed by conditioned media collection at 72 h for downstream functional and multiplexed analyses. **b**, Individual cytokine and chemokine measurements corresponding to Fig. 5a, quantified by multiplex electrochemiluminescence (MSD). Bar plots show levels of SASP-associated factors across treatment conditions. Significant p values are in red. **c**, Representative brightfield and eTox Red images of organoids exposed to conditioned media from control, IMF, IMF + BQ-788, or IMF + UPA conditions, corresponding to Fig. 5c. Insets show higher-magnification views. **d**, Time-course measurements of transepithelial electrical resistance (TEER) over 0–72 h, corresponding to Fig. 5d. Right, summary quantification at 72 h. Across assays, EDNRB antagonism consistently attenuates IMF-driven inflammatory and epithelial-disruptive phenotypes, with partial or context-dependent effects observed with JAK inhibition. *Statistics*: Results represent findings of conditioned media from independent donor-derived cultures; n = 2 unique IMF; 3-5 unique passages/subject. Bars denote mean ± S.E.M. Statistical comparisons were performed using one-way ANOVA or two-sided, unpaired t-tests, with exact *P* values shown in panels.

**Extended Data Figure 9.**
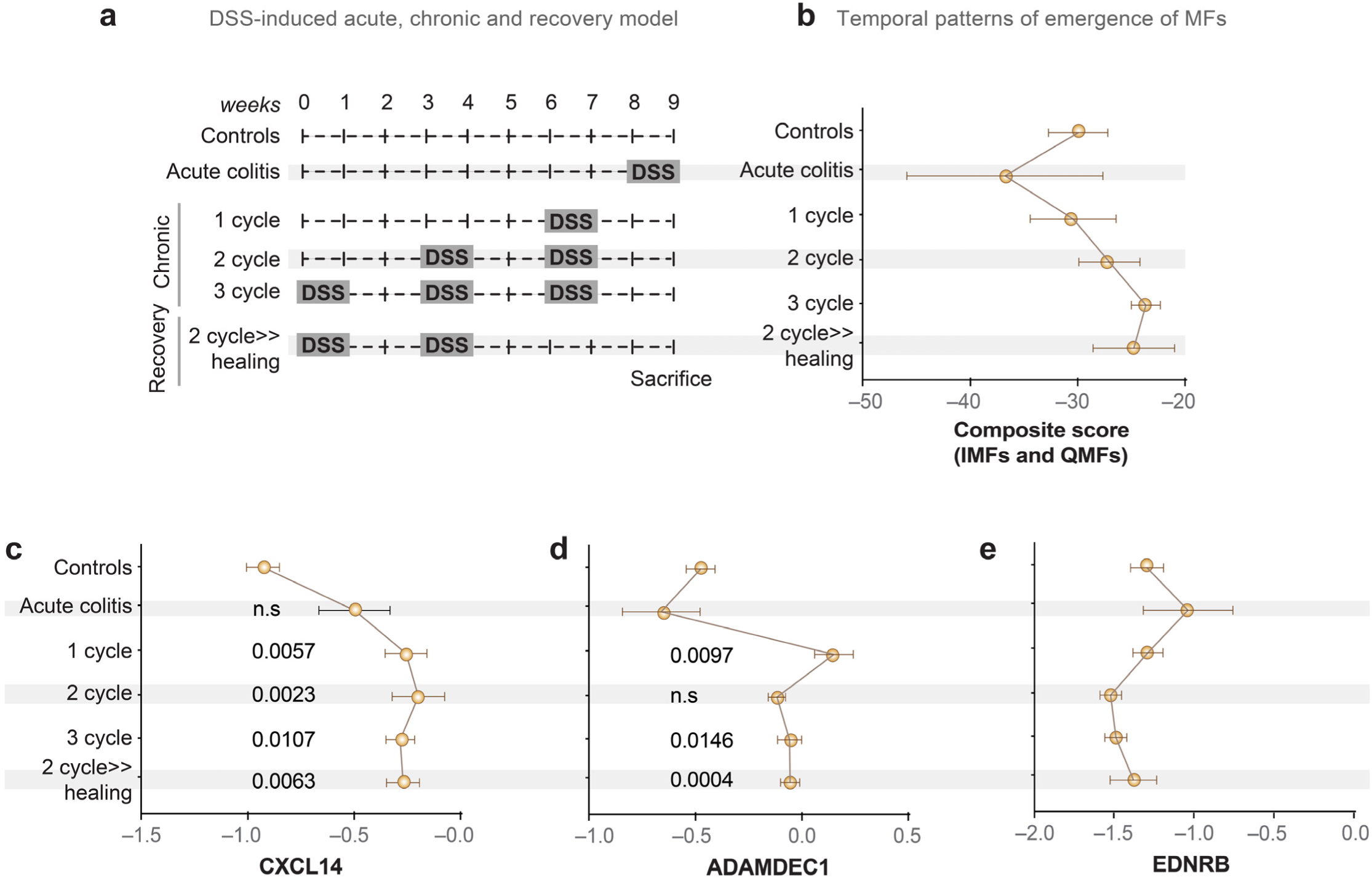
Temporal dynamics of stromal state emergence in DSS colitis inform therapeutic intervention windows. **a**, Schematic of DSS-induced acute, chronic, and recovery colitis models, as published before^79^. Mice were subjected to single or repeated DSS cycles to model progressive injury, followed by defined recovery phases prior to tissue harvest and bulk RNA sequencing (<u>GSE42768</u>). **b**, Composite stromal state score (IMF and QMF signatures) across disease stages, illustrating the temporal emergence and persistence of stromal activation during acute injury, chronic cycling, and recovery. **c–e**, Expression dynamics of representative stromal markers across disease progression: **c**, CXCL14 (IMF-associated), **d**, ADAMDEC1 (stromal activation marker), and **e**, EDNRB (therapeutic target). Data are shown across control, acute colitis, and increasing DSS cycles, including recovery conditions. Points indicate mean values with error bars (± S.E.M.), with statistical comparisons annotated. Significance was determined by two-tailed, unpaired t-test (all comparisons are to ‘control’ tissues). These data demonstrate that inflammatory stromal programs emerge early, intensify with repeated injury, and persist despite recovery, while EDNRB expression peaks during acute colitis, but remains low during post-colitic stages (2 weeks after injury) regardless of repeated injury. Together, these findings provide a rationale for timing EDNRB-targeted intervention during active inflammation and early chronic remodeling phases.

**Extended Data Figure 10.**
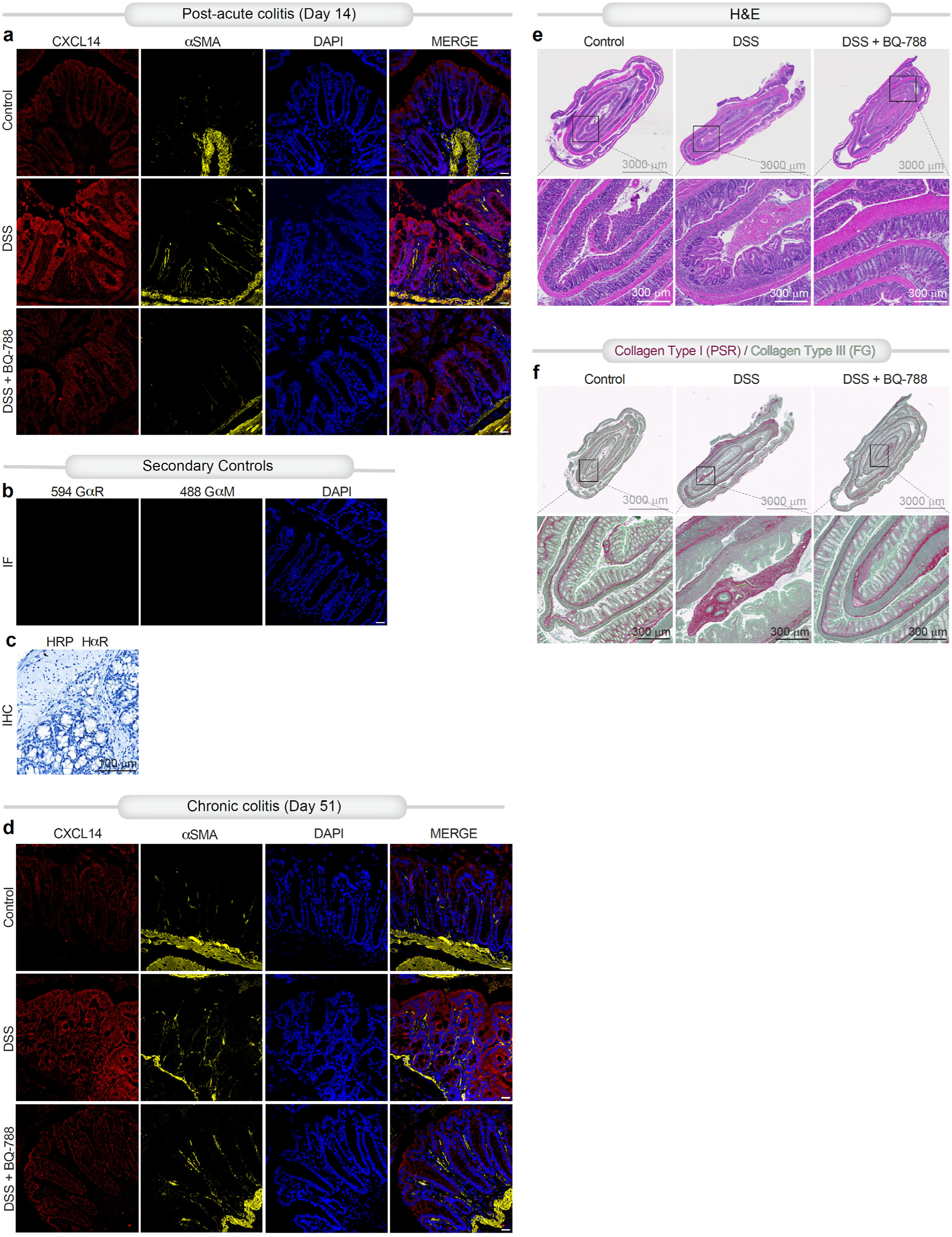
Tissue-level validation and controls for EDNRB-dependent stromal phenotypes in DSS colitis. **a, d**, Immunofluorescence staining of colonic tissues showing CXCL14 (IMF-associated; red), αSMA (stromal cells; yellow), and nuclei (DAPI; blue) in post-acute colitis (day 14; **a**) and chronic colitis (day 51; **d**) across control, DSS, and DSS + EDNRB antagonist (BQ-788) conditions. Representative fields illustrate expansion of CXCL14⁺ stromal compartments in DSS and attenuation upon EDNRB inhibition. Merged images highlight spatial colocalization within the lamina propria. Scale bars, 20 μm. **b**, Secondary antibody–only immunofluorescence controls demonstrating minimal background signal for goat anti-rabbit (GαR) and goat anti-mouse (GαM) antibodies, with DAPI nuclear counterstain. Scale bar, 20 μm. **c**, Immunohistochemistry (IHC) negative control (HRP-conjugated secondary antibody only) confirming specificity of staining and absence of nonspecific signal. Scale bar, 100 μm. **e**, Hematoxylin and eosin (H&E) staining of colonic tissue sections across experimental groups. Low-magnification images (*top*) show overall tissue architecture; boxed regions indicate areas shown at higher magnification (*bottom*). DSS induces epithelial distortion, inflammatory infiltration, and architectural disruption, which are partially restored by EDNRB antagonism. Scale bars, 3000 μm (top), 300 μm (bottom). **f**, Collagen deposition assessed by Picrosirius Red (PSR; collagen type I) and Fast Green (FG; collagen type III) staining. DSS induces increased collagen deposition and tissue remodeling, which are reduced upon EDNRB inhibition. Representative low- and high-magnification images are shown. Scale bars, 3000 μm (top), 300 μm (bottom). *Reproducibility*: Representative images are shown from independent biological replicates with multiple fields of view per condition. All staining and imaging were performed using identical acquisition and processing parameters across groups.

**Extended Data 11.**
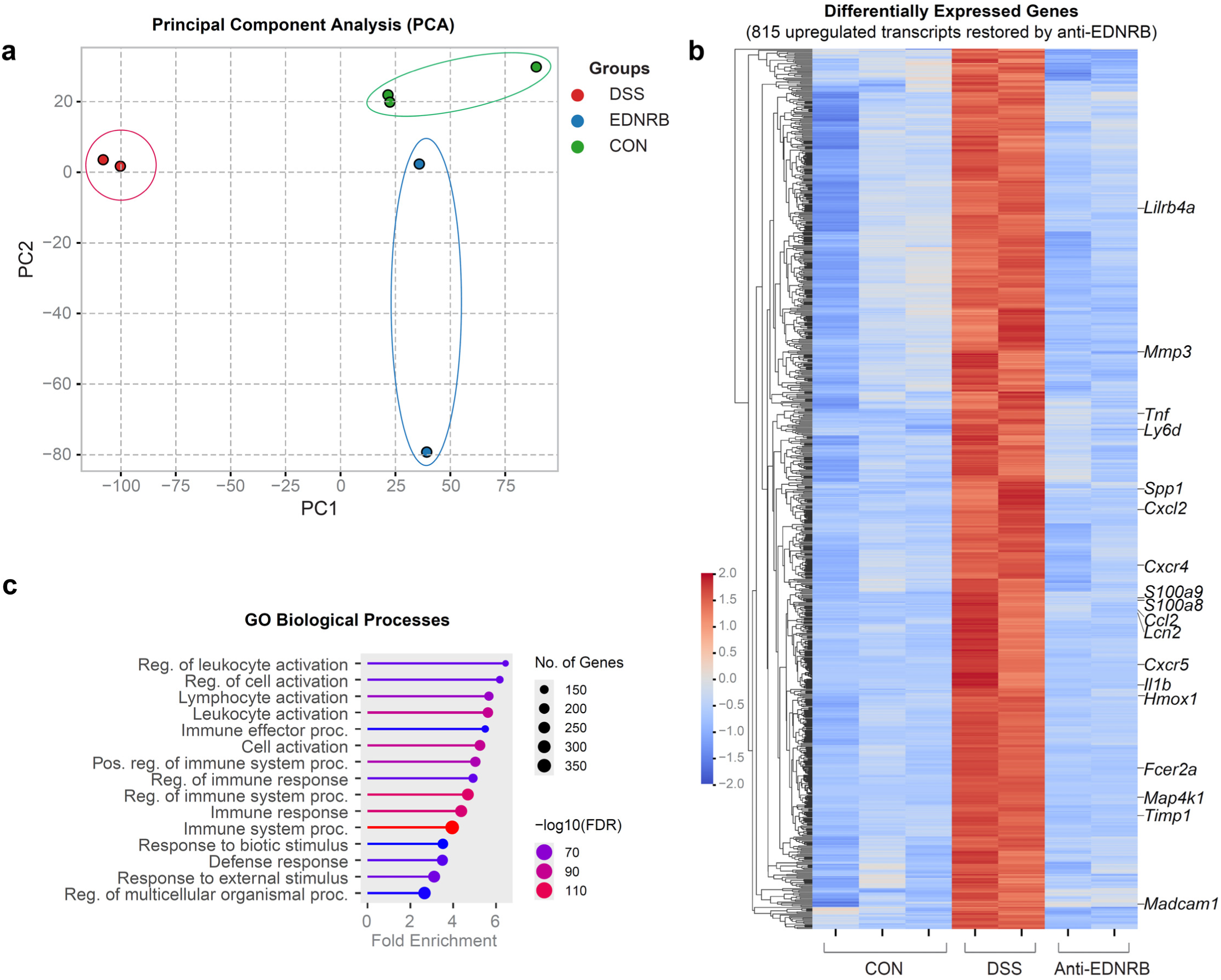
EDNRB antagonism reverses inflammatory transcriptional programs in DSS colitis. **a**, Principal component analysis (PCA) of bulk transcriptomic profiles from colonic tissue across control (CON), DSS-induced colitis (DSS), and EDNRB antagonist–treated (anti-EDNRB) conditions. Samples segregate by treatment group, with EDNRB antagonism partially restoring the DSS-induced transcriptional shift toward the control state. **b**, Heatmap of differentially expressed genes (DEGs) identified in DSS versus control tissue, highlighting 815 transcripts upregulated in DSS and restored toward baseline following EDNRB antagonism. Selected representative genes associated with inflammatory signaling, matrix remodeling, and immune activation are annotated. **c**, Gene Ontology (GO) enrichment analysis of DSS-upregulated genes reversed by EDNRB antagonism. Enriched pathways include leukocyte activation, lymphocyte activation, immune effector processes, and responses to external and biotic stimuli, indicating suppression of inflammatory and immune activation programs upon treatment. Together, these analyses demonstrate that EDNRB antagonism reprograms disease-associated transcriptional networks, reversing core inflammatory pathways induced during colitis. *Statistics*: Transcriptomic analyses were performed on independent biological samples as indicated. Differential expression and enrichment analyses were conducted using standard pipelines with multiple testing correction (FDR), with significance thresholds as indicated in panels.

**Extended Data 12.**
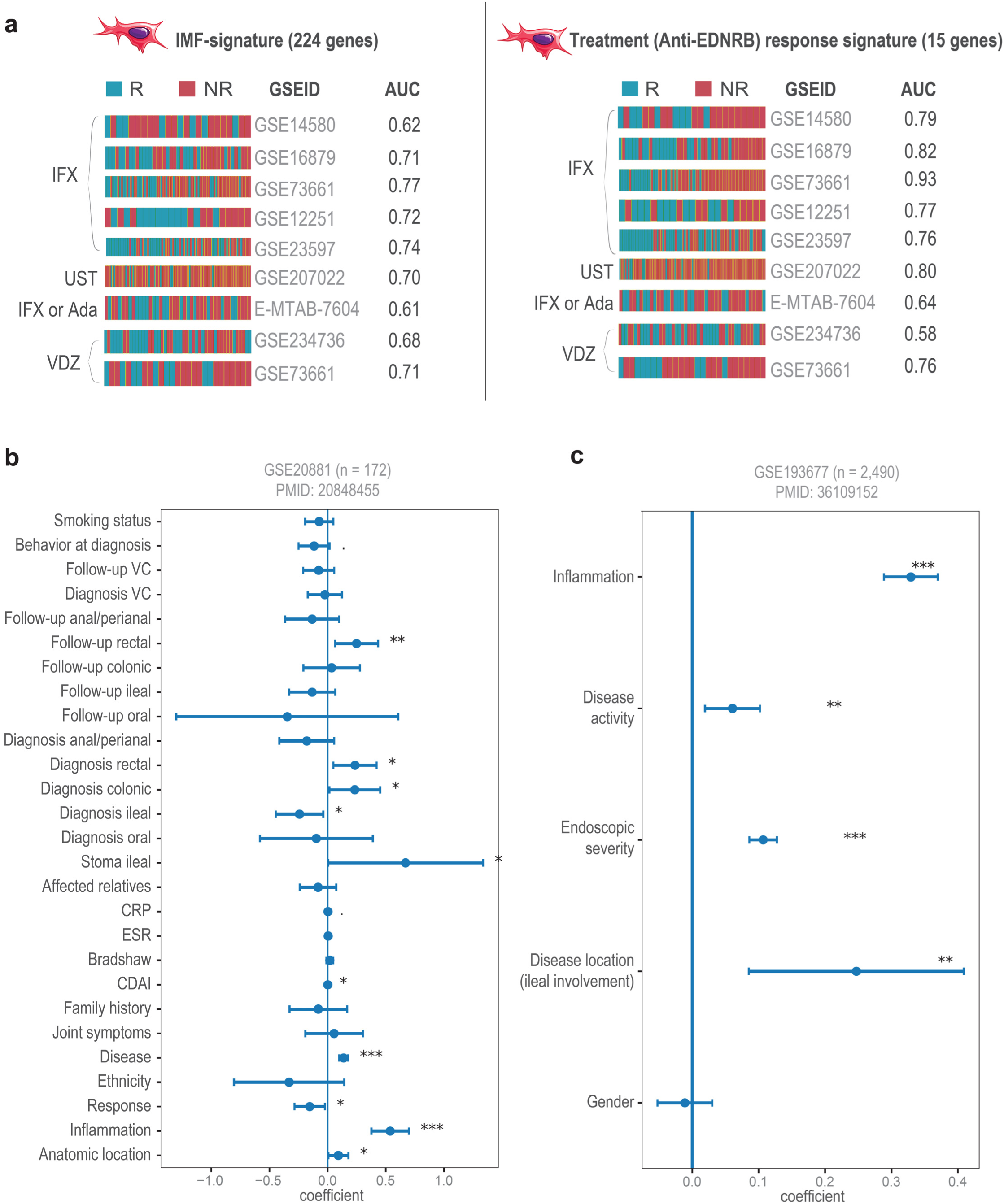
Benchmarking and clinical associations of stromal state–derived signatures. **a**, Comparative performance of the full IMF state–specific program (224 genes; left) versus the refined EDNRB response signature (15 genes; right) across independent transcriptomic datasets of biologic-treated IBD cohorts (IFX, infliximab; UST, ustekinumab; VDZ, vedolizumab). Heat strips indicate responder (R) and non-responder (NR) classifications, with corresponding ROC–AUC values shown for each dataset. The 15-gene EDNRB response signature demonstrates improved or comparable predictive performance relative to the broader IMF program. **b–c**, Association of stromal signatures with clinical disease features in independent cross-sectional cohorts. **b**, Univariate regression analysis (GSE20881; *n* = 172) showing associations between stromal signature scores and clinical variables, including disease location, inflammatory burden, and treatment response. **c**, Independent validation (GSE193677; *n* = 2,490) demonstrating significant associations of the stromal response program with inflammation, disease activity, endoscopic severity, and ileal involvement. Points represent regression coefficients with 95% confidence intervals; significance is indicated in panels. P < 0.05 (*); <0.01 (**); <0.001 (***). Multivariate analyses are presented in Fig 6h**-i**. Together, these analyses demonstrate that the EDNRB-derived stromal response signature is robust across cohorts, outperforms broader stromal gene sets, and captures clinically meaningful dimensions of disease severity and treatment response.

## Supplementary information

**Extended Data Figures (Extended Data Figures 1-12)**

**Extended Data Figure 1.** Isolation and characterization of stromal cell populations and pathway enrichment analyses, related to Fig. 1.

**Extended Data Figure 2.** Donor tissue validation of IMF and QMF stromal states across inflammatory contexts in IMF-predominant patients, related to Fig. 2.

**Extended Data Figure 3**. Donor tissue validation of IMF and QMF stromal states across inflammatory contexts in QMF-predominant patients, related to Fig. 2.

**Extended Data Figure 4**. Donor-tissue benchmarking of IMF and QMF stromal states in culture, related to Fig. 2.

**Extended Data Figure 5.** Metabolic and extracellular matrix features of QMF and IMF stromal states, related to Fig. 3.

**Extended Data Figure 6.** Cross-tissue validation of myofibroblast states and extended analysis of epithelial barrier dysfunction, related to Fig. 4.

**Extended Data Figure 7.** Stromal EDNRB expression is upregulated in IMF colonic mucosal tissue sections, related to Fig. 4.

**Extended Data Figure 8.** Pharmacologic modulation of IMF-derived paracrine programs in human NAMs, related to Fig. 5.

**Extended Data Figure 9.** Temporal dynamics of stromal state emergence in DSS colitis inform therapeutic intervention windows, related to Fig. 5.

**Extended Data Figure 10.** Tissue-level validation and controls for EDNRB-dependent stromal phenotypes in DSS colitis, related to Fig. 5.

**Extended Data 11.** EDNRB antagonism reverses inflammatory transcriptional programs in DSS colitis, related to Fig. 6.

**Extended Data 12.** Benchmarking and clinical associations of stromal state–derived signatures, related to Fig. 6.

**Supplementary Tables (Supplementary Table 1-3)**

**Supplementary Table 1**: Table of prior art.

**Supplementary Table 2**: Cross-cohort digital benchmarking of Inflammatory Myofibroblast (IMF) cultures in current work against fibroblast/myofibroblast subsets in inflammatory Bowel Disease across other single-cell transcriptomic studies, related to Fig. 2b and 6c.

**Supplementary Table 3**: List of chemicals/materials used in this study.

Supplementary Data (Supplementary Data 1-4)

**Supplementary Data 1**: Cohort characteristics at diversion and during follow-up, related to Fig. 1a-c, 2h-i, 6j-k.

**Supplementary Data 2**: List of differentially expressed 224 genes, used as IMF signature, related to Fig. 1d, 6a.

**Supplementary Data 3**: List of genes that track stromal trajectory states, related to Fig. 1k-l.

**Supplementary Data 4**: List of differentially expressed 815 genes, that are induced by DSS injury, but downregulated by EDNRB inhibitor in murine colon, related to Fig. 6a, Extended Data 12a-b.

