## Supplementary material for "A stromal-state dichotomy governs recalcitrance or remission in inflammatory bowel disease": SOM

**Affiliations:**
[1] Department of Cellular and Molecular Medicine, University of California La Jolla, CA, USA.
[2] UC San Diego HUMANOID™ Center of Research Excellence, La Jolla, CA, USA.
[3] Department of Medicine, University of California San Diego, La Jolla, CA, USA.
[4] Center for Precision Computational Systems Network (PreCSN), University of California San Diego, La Jolla, CA, USA.

[†]These authors contributed equally

[‡]**Senior corresponding authors**:

**Brigid S. Boland, M.D.;** Associate Professor, Department of Medicine, University of California San Diego; 9452 Medical Center Drive, 1W 503, La Jolla, CA 92093. **Phone**: (858) 246-2353: **Email:**

**Pradipta Ghosh, M.D.;** Professor, Departments of Medicine, and Cellular and Molecular Medicine, University of California San Diego; 9500 Gilman Drive (MC 0651), George E. Palade Building, Rm 231, 239; La Jolla, CA 92093. Phone: 858-822-7633: Fax: 858-822-7636:

### CATALOG OF SUPPLEMENTARY MATERIALS

Supplemental Tables 1-3

**Table 1.** Table of prior art showcasing how current effort compares against other studies on intestinal stromal states, with an emphasis on studies that included inflammatory bowel diseases (IBD).

| **Author** (Year) | **Study design** (prospective vs retrospective) | **Disease ('n')** (UC; CD; subtypes) | **Multiplexed phenotyping** (homotypic, heterotypic) | **Biobanking** (expandable and cryo-preservable) | **Benchmarking** (donor-tissue; cross-cohort) | **Demonstrable use in therapeutic screens** | **Quantifiable biomarker for translation** | **Gene signature** (digital biomarker) | **Systems modeling of clinical outcomes** | **Take Home Message / Study Vision** |
|---|---|---|---|---|---|---|---|---|---|---|
| Current study (2026) | Prospective; two independent cohorts | UC, n=9<br>CD, n=23<br>Healthy, n=2<br>(total n = 34) | MF monocultures. MF-PDO co-cultures MF-monocyte co-cultures | Yes | Yes (Immunocytochemistry on donor tissues).<br>Yes (digital signatures across other biobanks) | Yes (includes 1 novel target, EDNRB and novel insights into the efficacy of JAKinibs as second/third-line therapy for recalcitrant disease) | Yes (secreted, and *in situ* demonstration of disease-defining biomarker, CXCL14) | Yes; **224 genes** (IMF); **15 genes** EDNRB response signature | Yes, prospective follow up ~5 y from tissue diversion | IBD stromal states can be prospectively captured, propagated, and therapeutically reprogrammed |
| McGregor et al. (2025) PMID: 41224999 | Retrospective | CD and non-CD associated fistulas<br><br>Total n=48 | MF monocultures. Controls and overexpressed *OSR1* and *TWIST1* fibroblasts | Not clear if fibroblasts were cryopreserved. Fibroblasts were expanded | Single cell RNAseq, spatial tissue profiling (transcriptomics) colon biopsies, full thickness tissue | No | No | Yes, 18 genes | No | First catalog of fistula-associated epithelial, immune and stromal cell states linked to establishment of tunnelling anatomy. |
| Kong et al. (2025) PMID: 40562913 | Retrospective | CD, n=21 | No | No | Single-cell and spatial transcriptomics on donor tissue | No | No | Yes; 405 genes | No | Key transcriptomic and cellular networks in stricturing CD. |
| Mukherjee et al. (2023) PMID: 37507073 | Retrospective | CD, n=13 | MF monocultures | Cultured isolated MFs | scRNAseq on full thickness CD resections | Yes (includes 1 novel target – CDH11) | No | Yes; 12 genes | No | First scRNAseq atlas of strictured bowel. |
| Kong et al. (2023) PMID: 36720220 | Retrospective | CD, n=46 | No | No | scRNAseq on TI and colon CD samples | No | No | Yes, 3 genes | No | Inventory of cell-type- and organ-specific differences in CD and potential directions for therapeutic development |
| Friedrich et al. (2021) PMID: 34675383 | Retrospective; three cohorts | UC+CD+IBDu n=31<br><br>UC, n=72<br><br>CD, n=25 | MF monocultures | Isolated stromal cells. Not clear if they were cryopreserved. | Single cell and bulk RNAseq, quantitative histopathology, in situ localization; colon biopsies, | No | No | Yes, 111 genes (Non-Response module); 38 genes (PDPN/THY + IAF); 48 | Identification of non-response associated stromal modules, nominates | The identification of distinct, localized, tissular pathotypes will aid precision targeting of current therapeutics. |

| | | | | | | | | | | |
|---|---|---|---|---|---|---|---|---|---|---|
| | | | | | full thickness tissue | | | genes (ABCA8+, Progenitor); 78 genes (PDGFRA+; Progenitor) | IL-1 signaling blockade in ulcerating disease | |
| Thomas et al. (2024) [PMID: 39438660] | Prospective; TAURUS-IBD Study | CD, n=16 UC, n=22 | No | No | Single cell RNAseq, spatial transcriptomics colon mucosal biopsies | No | No | Yes, 75 IBD gene programs (pFP11, pFP01 represent stromal specific programs) | No | Authors describe a therapeutic atlas for CD and UC following anti-TNF treatment. |
| Gao et al. (2024) [PMID: 39303725] | Retrospective Pooled Analysis of Multiple Studies | 47,572 Colon Fibroblast Cells (Various Diseases, i.e. CRC, IBD, Healthy) | MF monocultures and co-cultures with T-cells or recombinant chemokines/ cytokines | MFs were isolated. Not clear if they were cryopreserved. | scRNAseq on 517 human samples, spanning 11 tissue types and pathologies | No | No | Yes, 264: C2 - ADAMDEC+ Lamina Propria Fib<br><br>229: C3 - COL15A1+ Progenitor-like Fib<br><br>203: C6 - ADHB1+ Alveolar Fib<br><br>337: C15 - SOX6+ Epithelial-crypt Fib | No | This study advances our understanding of fibroblast biology and suggests potential therapeutic strategies for targeting specific fibroblast subsets in cancer treatment. |
| Kinchen et al (2018) [PMID: 30270042] | Retrospective | UC, n=5 | Yes | Isolated stromal cells. Not clear if they were cryopreserved. | scRNAseq on UC mucosal colon biopsies | No | No | Yes; 280 genes marking the colonic crypt niche mesenchymal cell expressing SOX6 and Wnts (S2 Signature) | No | Authors identify a $SOX6^{+}F3^{+}$ WNT-producing mesenchymal niche near epithelial crypts and show how its remodeling drives inflammation and barrier dysfunction in IBD. |
| Pelka et al. (2021) [PMID: 34450029] | Retrospective | CRC, n=62 | Cytokine stimulation of monocultures | Yes; n=2 | scRNAseq on primary colorectal cancer tumor tissue | No | No | Yes, 150 genes that identify BMP-producing Fibroblasts | No | Authors reveal the logic underlying spatially organized immune-malignant cell networks. |

| | | | | | | | | | | |
|---|---|---|---|---|---|---|---|---|---|---|
| Hattangady et al. (2024) PMID 38385965 | Retrospective | Healthy colon fibroblasts, n=7 | MF monocultures | MFs were isolated and passaged. Not clear if they were cryopreserved. | Primary-derived colon fibroblasts Rx'd with SASP-inducing agent for transcriptome profiling | No | No | Yes, 306 genes representing core senescence profile in senescent primary human colon fibroblasts | No | Authors generate a SASP atlas of human colon fibroblasts |
| Yim et al. (2018) PMID 30589872 | Retrospective | CD, n=18 | No | MFs were isolated and passaged | Methylome and transcriptome from primary derived fibroblasts of CD non-inflamed, inflamed, stenotic mucosal tissue | No | No | Yes, 76 differentially methylated and expressed genes in fibroblasts isolated from stenotic (STEN) vs non-inflamed (NINF) CD tissue | No | Authors provide evidence that the methylome and the transcriptome are systematically dysregulated in stenosis-associated fibroblasts. |
| Otte et al. (2003) PMID: 12806620 | Retrospective | Healthy, CD; Cultured and primary-derived fibroblasts (old study) | MF Monocultures | MFs were isolated and passaged | TLR identification from cultured and primary-derived fibroblast study | No | No | Yes, 11 genes reflecting intestinal innate immune receptors | No | Bacterial components directly activate intestinal myofibroblasts expressing TLRs and these cells make have innate immune functions. |
| Chen et al. (2024) PMID 38302662 | Retrospective | Healthy, CD; n=120 | No | No | FFPE, qPCR | No | No | Yes, 14 gene signature with potential prognostic importance for strictures. | No | Findings support the importance of colonic gene expression in predicting outcomes for pediatric CD. |

1

**Table 2**: Cross-cohort digital benchmarking of Inflammatory Myofibroblast (IMF) cultures in current work against fibroblast/myofibroblast subsets in inflammatory Bowel Disease across other single-cell transcriptomic studies. p values denote the statistical comparisons of composite gene score distributions between QMF v IMF.

| Novel Class | Markers | Digital Biomarkers (Gene signatures) | Proposed Function | Tissue Location (CO: Colon, IL: Ileum, PA: Perianal) | Full Thickness vs. Biopsy | Disease: CD/UC Phenotype | Reference | P-Value |
|---|---|---|---|---|---|---|---|---|
| **Inflammatory Myofibroblast (IMF)-associated 224 genes** | CXCL14+, EDNRB+ | See **Datasheet 2** | Recalcitrant IBD (i.e., difficult to treat diseases not amenable to existing biologics) | Colon | Biopsies | CD, UC | Tindle et al, 2026 (This work) | 3.5e-14 |
| **$THY1^+FAP^+PDPN^+$ fibroblast**<br>PMID: 39438660 | THY1, PDPN, FAP, WNT2B, POSTN, CD74, | FTH1, SOD2, PDPN, NNMT, LUM, TIMP1, RARRES2, CXCL1, PHLDA1, COL3A1, PLAU, COL6A3, CHI3L1, COL1A1, TNFAIP6, DCN, PDGFRA, PDGFRB, ABCA8, WNT2B, FOS, SOX6, POSTN, CD74, C3, TNC, NRG1, LOCL2 | Anti-TNF non-remission | IL, CO | Biopsies | CD, UC | Thomas et al. 2024 | 6.51e-06 |
| **$MMP^+/WNT5A^+$ Fibroblasts**<br>PMID: 37507073 | MMP1, MMP3, WNT5A | SOD2, IER3, COL1A2, COL3A1, COL5A1, MMP1, MMP3, FTH1, MTCO2, MT-CO1, CD11, PHLDA1, CCN2, IL1R1, CD11, PLAU, CHI3L1, WNT2, IL24 | Intercellular interactions – Principal signal senders in fibrosis | IL | Full thickness | CD<br><br>Stricturing | Mukherjee et al. 2023* | 0.001 |

| | | | | | | | | |
|---|---|---|---|---|---|---|---|---|
| **S4** PMID: 30270042 | CCL19, CCL21, TNFSF14, CD74 | CCL21, CXCL13, TNFSF11, CD24, OLFM3, PTGDS, JAK3, PSMD2, IL32, IL33, CXCL2, CXCL3, TNFSF14, FDCSP, PDPN, C3 | Pro-inflammatory, immunomodulatory | CO | Biopsies | UC | Kinchen et al. 2018 | 0.003 |
| **Inflammation-Associated Fibroblasts (IAFs) – IL13RA2/IL11⁺** PMID: 31348891 | FAP, TWIST1, IL13RA2, IL11 | IL11, IL24, IL13RA2, WNT2B, WNT5A, WNT5B, FAP, TWIST1, WNT2, PLAU, CHI3L1, MMP3, TNFSF11, MMP10, IL1R1, OSMR, STRA6, IL1B, GOS2, IL8, IL10, CCL11, TNFRSF11B, NOTCH2NL, DKK3, CXCL12, BMP4, BMP5 | Mediate anti-TNF drug resistance, cell-to-cell communication, pro-fibrotic | CO | Biopsies | UC | Smillie et al. 2019 | 2.2e-05 |
| **Activated fibroblasts (Martin et al)** PMID: 31474370 | CD90, PDPN,CTHRC1, CHI3L1, PDGFRA, MMP2 | CXCL13, CXCL6, CXCL16, CXCL1, CXCL2, CXCL5, CXCL8, CCL2, CCL7, IL6, IL11 | Pro-inflammatory - neutrophil and monocyte recruitment, Pro-fibrotic | IL | Biopsies | CD | Martin et. al 2019 | 1.6e-04 |
| **Inflammatory fibroblasts IL11⁺CHI3L1⁺** PMID: 36720220 | CHI3L1, IL11, MMP3, MMP1, TNC | SOD2, TIMP1, COL6A3, TGFBI, CHI3L1, PHLDA1, COL1A1, IGFBP5A, SPON2, TNC, PCOLCE, IER3, MMP3 | Tissue remodeling, profibrotic and immunomodulatory signaling | CO | Biopsies | CD<br><br>Stricturing | Kong et al. 2023 | 0.002 |

| | | | | | | | | |
|---|---|---|---|---|---|---|---|---|
| **FAP⁺ fibroblasts** PMID: 39042469 | FAP, CD82, TWIST1, POSTN | TWIST1, SIX2, PRRX2, MSX2, HIF1A, CD90, IL6, IL11, TGFB1, CXCL1, CXCL5, CXCL6, CCL2, CCL7, IL32, CXCL16, IL32, CXCL10 | Fibrosis, immune modulation and inflammatory cell recruitment | IL | Full thickness | CD<br>Stricturing | Ke et al. 2024* | 0.006 |
| **IAF** PMID: 31730855 | IL1B, IL6 | TWIST1, PRRX1, TNC, COL1A2, F3, APOE, CXCL1, ADAMDEC1, MMP3, IL11, CXCL5, WNT5A, BMP1, IL1B, IL24, IL6 | Inflammatory signaling, stress adaptation, extracellular matrix organization | CO | Biopsies | UC (Paediatric) | Huang et al. 2019* | 0.006 |
| **PDPN⁺ Inflammatory/Activated Fibroblasts** PMID: 39169060 | PDPN, MMP1, SOD2 | CHI3L1, MMP1, PDPN, SOD2, COL7A1, MT2A, MMP3, SERPINE1, CEMIP, SCHLAP1, IGHG1, IL11, IGKC, CXCL5, IGLC3, CXCL1 | Central regulator of immune-stromal interactions | IL | Full thickness | CD | Li et al. 2024 | 0.008 |
| **CXCL14/ADAMDEC1⁺** PMID: 37507073 | ADAMDEC1, CXCL14, CCL11 | CXCL14, APOE, FABP5, IGFBP7, COL3A1, COL18A1, IL1R1, MT-CO2, MT-CO1, CDH11 | Intercellular interactions – Central signal senders in fibrosis<br>Tissue remodeling – increased ECM production | IL | Full thickness | CD<br>Stricturing | Mukherjee et al. 2023* | 0.042 |
| **Fistula-associated stromal (FAS) fibroblasts** PMID: 41224999 | WNT5A, TWIST1, TWIST2, PRRX1, PRRX2 and OSR2 | INHBA, PRRX2, IL24, CXCL5, CXCL13, IL11, TWIST1, CHI3L1, TWIST2, PRRX1, POSTN, CCL19, RUNX2, MMP3, MMP1, COL7A1, WNTF5A, F3 | FAS populations associate with distinct collagen structures, displaying properties of collagen deposition and fibrosis, detected in fistula and base of non-penetrating ulcers. | Mixed | Biopsies | CD | McGregor et al. 2025 | 0.032 |

| | | | | | | | | |
|---|---|---|---|---|---|---|---|---|
| **CXCL10⁺CCL19⁺** PMID: 35649411 | CCL19, CCL2, CCL13, CD74, HLA-DRA, VCAM1, CCL5 | CCL19, CXCL19, CD74, HLA-DRA, RBP5, CCL21, CXCL10, IRF8M, OASL, CXCL11, IL32, TNFSF13B, HLA-DRB1 | Immune cell interaction, antigen presentation, T cell activation, strong interferon response, pro-inflammatory signaling. | CO | Biopsies | UC | Korsunsky et al. 2022* | 0.038 |
| **CXCL14/F3/PDGFRA⁺** PMID: 37507073 | F3, CXCL14, PDGFRA, POSTN | MT-CO2, MT-CO1, IGFBP3, PHGR1, CDH11, PLAU, ILIR1 | Intercellular interactions – Central signal senders in fibrosis<br>Tissue remodeling – increased ECM production | IL | Full thickness | CD<br><br>Stricturing | Mukherjee et al. 2023* | 0.16 |
| **Inflammatory fibroblasts CHI3L1hi** PMID: 38663404 | CHI3L1, MMP1 | MMP1, MMP3, MMP13, MT1E, MT1M, MT1X, FOSB, EGR1, GLUL, FOS, CHI3L2, CHI3L1, CXCL13, CLU, CA12, NFKBIZ, SOD2 | Pro-fibrotic, cellular movement, invasion and migration, leucocyte migration | Perianal fistula | Fistula tract biopsy | CD<br><br>Perianal fistula | Levantovsky et al. 2024 | 0.14 |
| **Activated fibroblasts (Mennillo et al)** PMID: 38374043 | CD10, TIMP1, IL1R1, CD44, CXCL14, MMP1, MMP3 | TIMP1, MMP1, MMP3, AREG, TMEM158, TNFRSF11B, PDGFRA | Increased signature in vedolizumab nonresponders | CO | Biopsies | UC | Mennillo et al. 2024* | 0.16 |
| **LUM⁺ fibroblasts (Fibroblast cluster 9 (C9) and 12 (C12))** PMID: 38685679 | LUM<br>C9: DCN<br>C12: JUN | COL4A1, COL4A2, COL15A1, COL6A3, COL18A1, ADAMDEC1, LAMB1, GREM1 | C9 & C12: Pro-fibrotic, ECM and tissue structure organization, C9: WNT signaling regulation | IL | Full thickness | CD<br><br>Stricturing | Humphreys et al. 2024* | 0.5 |
| **S4 Fibroblast (FLC)** PMID: 33290721 | CCL19, FDCSP | COL1A1, COL6A1, FOXL1, ADAMDEC1, ZEB2, TAGLN, APOE | Pro-inflammatory signaling, epithelial-stromal crosstalk, WNT-mediated TA cell signaling, TNF-related epithelial apoptosis, | IL | Biopsies | CD (Paediatric) | Elmentaite et al. 2020 | 0.2 |

| | | | | | | | | |
|---|---|---|---|---|---|---|---|---|
| | | | altered epithelial cell composition | | | | | |
| **TWIST1⁺FAP⁺ fibroblasts** PMID: 39024569 | TWIST1, FAP, CHI3L1, CTHRC1, IGFBP4, GREM1, CTSK, THBS2, C3, FBLN2 | COL1A1, COL3A1, FAP, TWIST1, CHI3L1, NT5E, GREM1, FNDC1 | Tissue remodeling – ECM and structural organization | IL | Full thickness | CD<br>Stricturing | Zhang et al. 2024* | 0.012 |
| **SPARC⁺COL3A1⁺** PMID: 35649411 | COL11A1, SPARC, LRRC15, MMP13, MMP11, COL3A1 COL1A1, TGFB1 | CTHRC1, COL1A1, KIF28B, POSTN, ADAM12, COL3A1, MMP13, SPARC, CTHRC1, KIF26B, COL11A1, CRABP2, LRRC15, MMP11, FAP, COL5A1, TGFB1 | ECM remodeling, tissue development, matrix disassembly, structural support. | CO | Biopsies | UC | Korsunsky et al. 2022* | 0.004 |
| **Inflammatory fibroblasts** PMID: 34675383 | PDPN, THY1, FAP | MMP13, MMP3, MMP1, CRABP2, CXCL5, DPT, CXCL6, TNFAIP6, IL11, PTGFR, FDCSP, CA12, OGN, CHI3L1, ADAM12, PDPN | IL-1R dependent neutrophil-chemoattraction | IL, CO | Biopsies<br>Full thickness | UC, CD | Friedrich et al. 2021* | 0.092 |

**Table 3**. Table of key resources used in this study.

| Reagent or Resource | Source | Identifier |
|---|---|---|
| **Antibodies** | | |
| Anti-Collagen I antibody [EPR24331-53] | Abcam | Cat# ab270993 |
| Anti-Ki67 antibody [SP6] | Abcam | ab16667 |
| P-Histone H2A.X (Ser 139) | Santa Cruz | sc-517348 |
| CD90 Monoclonal Antibody (5E10), PE - A15794 | Invitrogen | A15794 |
| Alpha-Smooth Muscle Actin Monoclonal Antibody (1A4), eFluor™ 660 (Flow cytometry) | Invitrogen | 50-112-4530 |
| Alpha-Smooth Muscle Actin Monoclonal Antibody (1A4), eBioscience (IF) | Invitrogen | 14-9760-82 |
| Alpha-Smooth Muscle Actin Monoclonal Antibody (EPR5368) (IF) | Abcam | ab124964 |
| Percp/cy5.5 anti-human CD274 (B7-H1, PD-L1) [29E.2A3] | BioLegend | 329738 |
| CXCL14 Polyclonal antibody | Proteintech | Cat# 10468-1-AP |
| CD36 Monoclonal antibody | Invitrogen | Cat# TA500921 |
| EDNRB | Proteintech | Cat# 20964-1-AP |
| CD45 Alexa Fluor 488 | Biolegend | Cat# 36853 |
| Goat Anti-Rabbit Alexa Fluor 488 Antibody | Abcam | Cat# 150077<br>RRID: AB_2630356 |
| Goat Anti-Mouse Alexa Fluor 488 Antibody (IF) | Invitrogen | A11001 |
| Goat Anti-Rabbit Alexa Fluor 594 Antibody (IF) | Invitrogen | A11012 |
| **Experimental Models: Organisms/ strains** | | |
| C57BL/6 Mice | Jackson Laboratory | Strain #:000664<br>RRID:IMSR_JAX:000664 |
| Adherent-Invasive *E. coli* LF82 | Arlette Darfeuille-Michaud | NA |
| **Chemicals, peptides, and recombinant proteins** | | |
| MEM | Corning | Cat# MT 10-010-CV |
| Collagenase Type I | Thermo Fisher | Cat# 17100017 |
| HBSS with Ca2+/Mg2+ | Thermo Fisher | Cat# 14-025-092 |
| HBSS without Ca2+/Mg2+ | Thermo Fisher | Cat# 14-175-095 |
| MEM Non-Essential Amino Acids | Thermo Fisher | Cat# 11-140-050 |
| Intestigro™ [L-WRN conditioned media, for intestine and colon PDOs] | UC San Diego HUMANOID™ Center | Cat# HUM2019 |
| Tailor-2-Gro™ [L-WRN conditioned media, base[P] | UC San Diego HUMANOID™ Center | Cat# HUM202420 |
| Seahorse XF DMEM assay medium pack | Agilent Technologies | Cat# 103680-100 |
| Sodium Pyruvate | Sigma-Aldrich | Cat# S8636 |
| Antibiotic Antimycotic | Sigma-Aldrich | Cat# A5955 |
| Bafilomycin A1, ≥90% (HPLC), From Streptomyces griseus | Sigma-Aldrich | Cat# B1793-10UG |
| Trypsin (2.5%) | Thermo Fisher | Cat# 15090046 |
| Zinc Formalin | Fisher Scientific | Cat# 23-313096 |
| Xylene | VWR | Cat# XX0060-4 |
| Hematoxylin | Sigma-Aldrich Inc | Cat# MHS1 |
| Ethanol | Koptec | Cat# UN1170 |

| | | |
|---|---|---|
| Sodium Citrate | Sigma-Aldrich | Cat# W302600 |
| DAB (10x) | Thermo Fisher | Cat# 1855920 |
| Stable Peroxidase substrate buffer (10x) | Thermo Fisher | Cat# 34062 |
| 3% Hydrogen Peroxide | Sigma | Cat# 245-07-3628 |
| Goat Serum | Vector Labs | Cat# S-1012-50 |
| Horse Serum | Vector Labs | Cat# S-2012-50 |
| Paraformaldehyde 16% Solution, EM Grade | Electron Microscopy Sciences | Cat# 15710 |
| 100% Methanol | Supelco | Cat# MX0485 |
| Glycine | Fisher Scientific | Cat# BP381-5 |
| Bovine Serum Albumin | Sigma-Aldrich | Cat# A9647-100G |
| Triton-X 100 | Sigma-Aldrich | Cat# X100-500ML |
| Prolong Glass | Invitrogen | Cat# P36984 |
| Nail Polish (Rapid Dry) | Electron Microscopy Sciences | Cat# 72180 |
| Gill Modified Hematoxylin (Solution II) | Millipore Sigma | Cat# 65066-85 |
| TrypLE Select | Thermo Scientific | Cat# 12563-011 |
| Advanced DMEM/F-12 | Thermo Scientific | Cat# 12634-010 |
| Glutamax | Thermo Scientific | Cat# 35050-061 |
| Penicillin-Streptomycin | Thermo Scientific | Cat# 15140-122 |
| Matrigel | Corning | Cat# 354234 |
| DPBS | Thermo Scientific | Cat# 14190-144 |
| Ultrapure Water | Invitrogen | Cat# 10977-015 |
| EDTA | Thermo Scientific | Cat# AM9260G |
| Fetal Bovine Serum | Sigma-Aldrich | Cat# F2442-500ML |
| Cell Recovery Solution | Corning | Cat# 354253 |
| Sodium Azide | Fisher Scientific | Cat# S227I-100 |
| Cyto-Fast Fix/Perm Buffer Set | BioLegend | Cat# 426803 |
| Human TruStain FcX™ (Fc Receptor Blocking Solution) | BioLegend | Cat# 422302 |
| FITC-Dextran | Sigma-Aldrich | Cat# FD10S |
| Ethyl alcohol, pure | Sigma-Aldrich | Cat# E7023 |
| TRI Reagent | Zymo Research | Cat# R2050-1-200 |
| Dextran Sodium Sulfate | M.P. Biomedicals | Cat# 9011-18-1 |
| BQ-788 | MedChem Express | Cat# HY-15894A |
| Upadacitinib | MedChem Express | Cat# HY-19569 |
| **Critical commercial assays** | | |
| SPiDER-βGal | Dojindo | Cat# SG02-10 |
| Quick-RNA MicroPrep Kit | Zymo Research | Cat# R1051 |
| Quick-RNA MiniPrep Kit | Zymo Research | Cat# R1054 |
| Seahorse XF Mito Stress Test Kit | Agilent Technologies | Cat# 103015-100 |
| HRP Horse Anti-Rabbit IgG Polymer Detection Kit | Vector Laboratories | Cat# MP-7401 |
| Human CXCL14/BRAK DuoSet ELISA | R&D Systems | Cat# DY866 |
| **Software and algorithms** | | |

| | | |
|---|---|---|
| ImageJ/Fiji | ImageJ; https://imagej.net/Welcome | RRID:SCR_003070 |
| Illustrator | Adobe | https://www.adobe.com/products/illustrator.html |
| Python3 | Python | https://www.python.org |
| GraphPad Prism 9 | GraphPad Prism ; https://www.graphpad.com/scientific-software/prism/ | RRID:SCR_002798 |
| MSD® DISCOVERY WORKBENCH 4.0 | MSD | |
| LAS AF Software | LAS AF Software | www.leica-microsystems.com/products/microscope-software/p/leica-las-x-ls |
| FlowJo | Flow Jo V10, BD BioSciences | RRID:SCR_008520 |
| QuPath | QuPath v0.5 | https://qupath.github.io/ |
| **Other** | | |
| 12-well Tissue Culture Plate | CytoOne | Cat# CC7682-7512 |
| HTS Transwell®-96 Permeable Support with 0.4μm PET Membrane | Corning | Cat# 7369 |
| Cell Scraper | Millipore Sigma | Cat# C5981-100EA |
| Millicell EZ Slide 8-Well Chamber | Millipore Sigma | Cat# PEZGS0816 |
| Trypan Blue Stain | Invitrogen | Cat# T10282 |
| 70 um Cell Strainer | Thermo Fisher Scientific | Cat# 22-363-548 |
| Noyes Spring Scissors - Angled | Fine Science Tools | Cat# 15013-12 |
| DAPI | Invitrogen | Cat# D1306<br>RRID: AB_2629482 |
| Molecular Probes Alexa Fluor 594 Phalloidin | Thermo Fisher Scientific | Cat# A12381 |
| eAnnexin V Red | Agilent Technologies | Cat# 8711007 |
| eTox Red | Agilent Technologies | Cat# 8711009 |
| Countess II Automated Cell Counter | Thermo Fisher Scientific | AMQAX1000 |
| Automated TEER measurement system [REMS AutoSampler] | World Precision Instruments (WPI) | |
| Precellys® Tissue Homogenizer | Bertin Corp. | |
| MESO QuickPlex SQ 120 | MSD | |
| Leica TCS SPE Confocal | Leica Microsystems | TCS SPE |
| Power Pressure Cooker XL | Tristar Products | |
| Canon Rebel XS DLSR | Canon | |
| Light Microscope (brightfield images) | Carl Zeiss LLC | Axio Observer, Inverted; 491917-0001-000 |
| Spark 20M Multimode Microplate Reader | Tecan | |
| NanoQuant Infinite M200 | Tecan | |
| xCELLigence RTCA eSight | Agilent Technologies | |
| BioTek Cytation C10 Confocal Imaging Reader | Agilent Technologies | |
| Seahorse XF Pro Analyzer | Agilent Technologies | |
| NovoCyte Quanteon Flow Cytometer | Agilent Technologies | |
| Guava® easyCyte Benchtop Flow Cytometer | Millipore | Guava easyCyte 6 2L |
| MESO QuickPlex instrument | Mesoscale Discovery Inc. | SQ 120 |
